# Transcriptional repression and enhancer decommissioning silence cell cycle genes in postmitotic tissues

**DOI:** 10.1101/2024.05.06.592773

**Authors:** Elizabeth A. Fogarty, Elli M. Buchert, Yiqin Ma, Ava B. Nicely, Laura A. Buttitta

**Affiliations:** Molecular, Cellular and Developmental Biology, University of Michigan, Ann Arbor 48109

## Abstract

The mechanisms that maintain a non-cycling status in postmitotic tissues are not well understood. Many cell cycle genes have promoters and enhancers that remain accessible even when cells are terminally differentiated and in a non-cycling state, suggesting their repression must be maintained long term. In contrast, enhancer decommissioning has been observed for rate-limiting cell cycle genes in the *Drosophila* wing, a tissue where the cells die soon after eclosion, but it has been unclear if this also occurs in other contexts of terminal differentiation. In this study, we show that enhancer decommissioning also occurs at specific, rate-limiting cell cycle genes in the long-lived tissues of the *Drosophila* eye and brain, and we propose this loss of chromatin accessibility may help maintain a robust postmitotic state. We examined the decommissioned enhancers at specific rate-limiting cell cycle genes and show that they encode dynamic temporal and spatial expression patterns that include shared, as well as tissue-specific elements, resulting in broad gene expression with developmentally controlled temporal regulation. We extend our analysis to cell cycle gene expression and chromatin accessibility in the mammalian retina using a published dataset, and find that the principles of cell cycle gene regulation identified in terminally differentiating *Drosophila* tissues are conserved in the differentiating mammalian retina. We propose a robust, non-cycling status is maintained in long-lived postmitotic tissues through a combination of stable repression at most cell cycle gens, alongside enhancer decommissioning at specific rate-limiting cell cycle genes.

**Highlights:** In long-lived postmitotic *Drosophila* tissues, most cell cycle genes retain accessible chromatin despite persistent transcriptional downregulation.

Cell cycle genes with accessible enhancers remain activatable during terminal differentiation, suggesting their repression must be continuously maintained in the postmitotic state.

Long-lived postmitotic tissues decommission enhancers at specific, rate-limiting cell cycle genes in a developmentally regulated manner.

Genome-wide enhancer identification performed in cell culture misses many developmentally dynamic enhancers *in vivo*.

Decommissioned enhancers at cell cycle genes include shared and tissue-specific elements that in combination, result in broad gene expression with temporal regulation.

The principles of cell cycle gene regulation identified in *Drosophila* are conserved in the mammalian retina.

## Introduction

Transcriptional control for many genes can be simplified into two categories of gene regulation: housekeeping genes with broad, ubiquitous expression and developmentally dynamic genes with cell type- or temporally-specific expression patterns. Work in *Drosophila* cells has characterized fundamental differences between these modes of gene regulation and shown that housekeeping genes and developmentally dynamic genes have unique enhancer architectures and preferentially use different types of core promoters (Zabidi et al., 2015). For example, enhancers that activate housekeeping-type promoters are often shared across cell types and located near gene transcription start sites (TSS), while enhancers that pair with developmental-type promoters are more likely to exhibit cell type-specific activity and are predominantly found in introns or intergenic regions. In these high-throughput, genome-wide enhancer identification studies, cell cycle genes were characterized as enriched among housekeeping-type genes (Zabidi et al., 2015). This characterization is consistent with a number of other studies using genome-wide transcriptomic measurements across panels of tissues to classify housekeeping and tissue-specific genes (Chang et al., 2011; Dezso et al., 2008; Farre et al., 2007; Hsiao et al., 2001; Joshi et al., 2022; She et al., 2009). However, the designation of cell cycle genes as “housekeeping” belies the complex regulation of cell cycle genes *in vivo*, where these genes are subject to spatial and temporal developmental dynamics (Geng et al., 2001; Jones et al., 2000; Kakizuka et al., 1992; Lehman et al., 1999; Thacker et al., 2003). Most studies identifying housekeeping and tissue specific genes have utilized panels of only adult stage tissues and have therefore been limited to the resolution of spatial rather than temporal expression dynamics. Furthermore, some groups have measured gene expression breadth based on binary on/off designations in each tissue, without regard to variation in expression level across tissues or time points. Indeed, the large dynamic range of expression for cell cycle genes has been noted in at least one study that included both early developmental and adult tissues in an analysis of the mouse transcriptome, where it was observed that genes involved in mitosis and cytokinesis were generally lowly expressed across adult tissues but much more highly expressed in the embryo (Zhang et al., 2004). The complex developmental regulation of cell cycle genes is especially obvious in tissues undergoing cell cycle transitions that are temporally regulated and coordinated with terminal differentiation programs, as in mammalian muscle and neuronal lineages as well as *Drosophila* eye, wing and brain (Firth and Baker, 2005; Meserve and Duronio, 2017; Milan et al., 1996; Ruijtenberg and van den Heuvel, 2016; Schubiger and Palka, 1987; Siegrist et al., 2010).

When cells transition from a proliferating to a postmitotic state during development, cell cycle gene transcriptional control switches from activation to repression. This is thought to be largely mediated by the transcription factor complex E2F, which controls the expression of hundreds of cell cycle genes and can form an activating or repressive complex, based on its binding partners and the particular E2F subunit present in the complex (Fischer and Muller, 2017; Fischer et al., 2022). The E2F activator complex in *Drosophila* contains E2F1 with its dimerization partner DP, while the E2F repressive complex contains E2F1 or E2F2 complexed with the inhibitory Retinoblastoma family (Rbf) protein along with components of a conserved complex called DREAM, for dimerization partner (DP), retinoblastoma (RB)-like, E2F and MuvB (Korenjak et al., 2004). Rbf and/or DREAM serves a critical function in cell cycle gene silencing during cell cycle exit (Litovchick et al., 2007), but whether this complex continuously maintains cell cycle gene repression over the longer term in postmitotic tissues is unclear. Importantly, the E2F-Rb axis is highly evolutionarily conserved, with E2F and Rb homologs present across metazoans and even present in plants (Cross et al., 2011; Dewitte and Murray, 2003). Therefore, it is not surprising that the study of cell cycle regulation in *Drosophila* has provided important insights that apply across many other systems.

We previously showed that postmitotic cells in the *Drosophila* wing decommission enhancers at three specific rate-limiting cell cycle genes after cell cycle exit (Ma et al., 2019). These include: the G1-S cyclin Cyclin E (CycE), the activator E2F subunit E2F1, and the mitotic Cyclin/Cdk phosphatase that mediates mitotic entry Cdc25c (called String [Stg] in flies) (Neufeld et al., 1998). We suggested this enhancer decommissioning may contribute to the robustness of permanent cell cycle exit and is likely to be developmentally controlled, since even bypassing cell cycle exit and keeping cells in a cycling state could not prevent the closing of regulatory elements at these genes (Ma et al., 2019). The genomic loci for *cycE*, *stg* and *e2f1* are unique among *Drosophila* cell cycle genes in that they contain complex, modular cis-regulatory regions, making them similar in architecture to genes characterized as developmentally regulated (Andrade-Zapata and Baonza, 2014; Jones et al., 2000; Lehman et al., 1999; Lopes and Casares, 2015). Indeed, hundreds of *Drosophila* cell cycle genes exhibit a simple enhancer architecture similar to what has been described for housekeeping genes. We were surprised to find that these genes retain TSS-adjacent chromatin accessibility after transcriptional silencing, suggesting they are subject to continual, long-term repression after cell cycle exit. Thus, our emerging model for maintenance of the postmitotic state includes stable repression of hundreds of cell cycle genes - perhaps through long-term occupation of promoter-proximal regulatory regions by repressive E2F complexes - together with enhancer decommissioning at the few cell cycle genes with complex enhancer architecture. However, this model is largely based on chromatin accessibility and gene expression data from the *Drosophila* wing, which is a short-lived tissue where cells are destined to die by apoptosis shortly post eclosion (Link et al., 2007). One possibility is that short-lived tissues may employ alternative strategies to achieve a relatively short-term and/or less stringent repression of cell cycle genes and may not be representative of all tissues. Here, we address the question of whether postmitotic tissues that persist for the lifetime of the animal follow the same principles of cell cycle gene regulation. Further, we explore the evolutionary conservation of this model by analyzing chromatin accessibility and gene expression data for cell cycle genes in the developing mouse retina.

## Results and Discussion

We have previously shown in the developing *Drosophila* wing that most cell cycle gene loci have simple chromatin accessibility profiles, harboring a single region of open chromatin near the transcription start site (Ma et al., 2019). By assaying chromatin accessibility at time points before, during and after cell cycle exit, we observed that the chromatin accessibility at these genes is maintained after cell cycle exit. Indeed, cell cycle gene promoters remain accessible in the wing even at time points long after cell cycle exit has occurred, cell cycle gene transcripts are no longer expressed, and the tissue has initiated its terminal differentiation program. To address whether these findings represent a wing-specific phenomenon or are also representative of long-lived tissues, we chose to compare our findings in the wing to two tissues that persist in the adult fly, the eye and the brain. We selected the eye and brain for this comparison because these tissues are composed of diploid cells that persist throughout adulthood and their final cell cycles occur during metamorphosis with roughly similar timing to the wing. In the wing, epithelial cells undergo a final cell cycle between 10-24 hours after puparium formation (APF) (Milan et al., 1996; Schubiger and Palka, 1987). In the eye, cell cycle exit is much less temporally synchronous; a subset of photoreceptor and cone cells exit from the cell cycle during larval and early pupal stages as the spatiotemporal morphogenetic furrow sweeps across the eye (Firth and Baker, 2005; Wolff and Ready, 1991). This is followed by final cell cycles for pigment cell and bristle precursors in the pupa retina that complete by 24h APF (Buttitta et al., 2007; Meserve and Duronio, 2017). In the brain, neural stem cells (termed neuroblasts) give rise to intermediate cell types to ultimately produce multiple neuronal and glial subtypes (Maurange and Gould, 2005; Rajan et al., 2021). The majority of neuroblasts also exit from the cell cycle around 24h APF. Apart from eight central brain neuroblasts termed the “mushroom body” neuroblasts, the brain is nearly fully postmitotic after 24h APF (Homem et al., 2014; Siegrist et al., 2010). To confirm that the timing of cell cycle exit in these tissues corresponds with E2F-dependent transcriptional repression, we assayed for mitotic events via antibody staining against phosphorylated histone H3 as well as the silencing of cell cycle gene expression through a well-characterized E2F transcriptional activity reporter, *pcna*-GFP, based upon an E2F-regulated enhancer at the *proliferating cell nuclear antigen* locus (Thacker et al., 2003). In all three tissues, E2F transcriptional activity and mitoses are readily detected at 20h APF, largely silenced by 24h APF, and remained silenced at 44h APF (Fig. 1).

**Figure 1.**
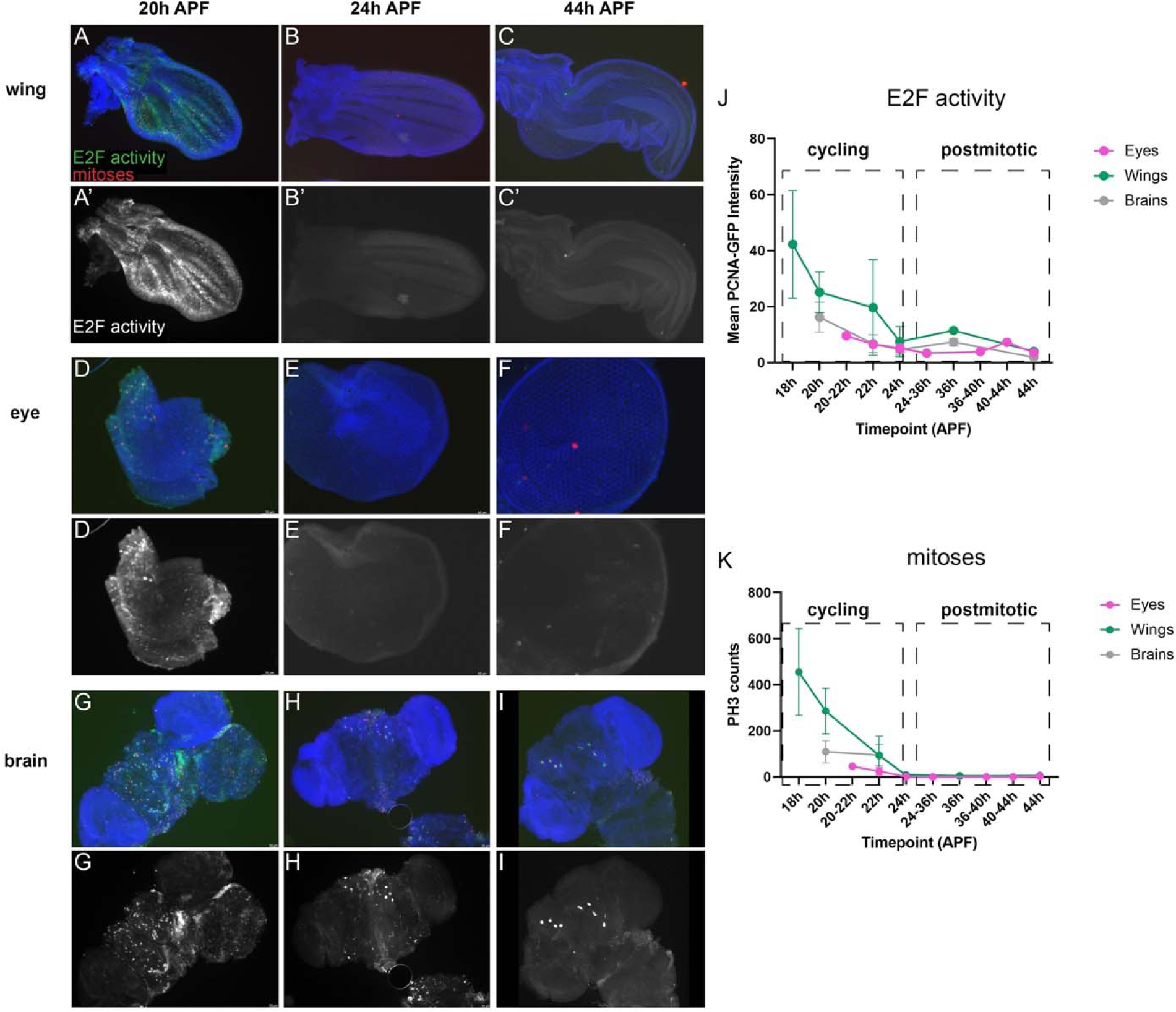
The timing of cell cycle exit in the *Drosophila* wing, eye and brain are similar. Wings (**A-C**), eyes (**D-F**) and brains (**G-I**) were dissected from staged pupa and stained for mitoses (anti-Phospho Histone H3, PH3) and E2F transcriptional activity (anti-GFP for PCNA-GFP reporter) at the timepoints indicated. Animals were collected as white pre-pupa (0h APF) and incubated at 25°C to the indicated timepoints. Yellow arrowheads indicate 4 of the 8 mushroom body neuroblasts that continue to cycle until 96h APF (Siegrist et al., 2010). J-K Quantifications of E2F transcriptional activity (normalized PCNA-GFP fluorescence) and PH3 across tissues at the indicated timepoints.

We next expanded upon our previous work by measuring chromatin accessibility and gene expression in wing, eye and brain at selected time points before, during and after cell cycle exit (Fig. 2A). Gene expression analysis by RNA-Seq confirmed that expression levels of cell cycle genes decrease in each tissue across this time course (Fig. 2B), in agreement with the cell cycle exit dynamics that are similar across tissues (Fig. 1). Consistent with what we previously reported in the wing, chromatin accessibility profiles as measured by ATAC-Seq reveal relatively simple regulatory architecture at most cell cycle genes in these tissues, with the majority of genes showing accessibility primarily at the TSS. This is supported by genomic distribution analysis of ATAC-Seq peaks, comparing all peaks genome-wide to those mapping closest to cell cycle genes. Peaks assigned to cell cycle genes are enriched at promoters and depleted at intronic and intergenic regions relative to the genome-wide distributions in all three tissues (Fig. 2C). Similarly, an analysis of the number of ATAC-Seq peaks annotated per gene revealed that cell cycle genes have fewer accessible regions than average, showing an enrichment of genes harboring a single peak and a depletion of genes with complex landscapes of five or more annotated peaks (Fig. 2D). Furthermore, despite the loss of transcript expression during and after cell cycle exit, we observed that chromatin accessibility is maintained at the peaks nearest cell cycle genes in each tissue - even at 44h APF when the tissues have been postmitotic for 20 hours and terminal differentiation is well underway (Fig. 2E). Taken together, these findings support the idea that what we previously observed in the wing, where the vast majority of cell cycle genes are subject to simple TSS-adjacent regulatory regions and retain accessibility after cell cycle exit, is a general principle of cell cycle gene regulation in *Drosophila* rather than a peculiarity of the short-lived wing cells.

**Figure 2.**
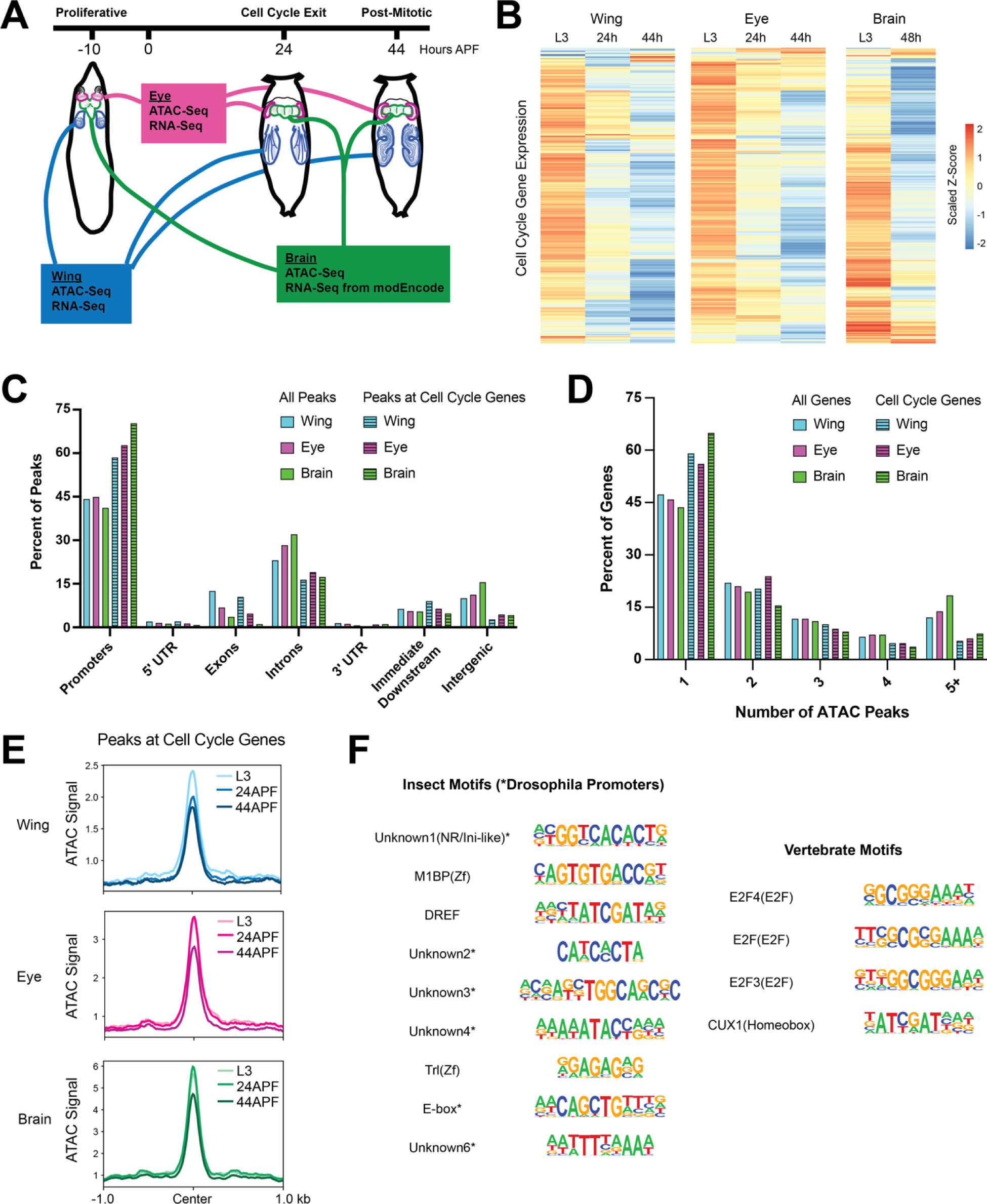
Most cell cycle genes have simpler than average regulatory architectures, are transcriptionally repressed after cell cycle exit, but retain chromatin accessibility in terminally differentiating fly tissues. (**A**) Wing, eye, and brain tissue was dissected from third instar larvae (L3, 10 hours prior to puparium formation) and from pupae at 24 hours or 44 hours after puparium formation (APF). ATAC-Seq and RNA-Seq datasets were generated from wing and eye tissues at all three time points. ATAC-Seq data was generated from brain tissue at all three time points. Publicly available RNA-Seq datasets from L3 larval brain and 2-day pupa brain were generated by modEncode. (**B**) Heatmap depicting the average RNA-Seq transcript expression values for 284 genes with annotated functions related to the cell cycle. Data are scaled by Z-score and hierarchically clustered. (**C**) Bar plot showing the genomic distributions of ATAC-Seq peaks from the wing, eye, and brain, either for all peaks (open bars) or only peaks assigned to cell cycle genes (striped bars) (**D**) Bar plot showing the distributions of genes binned by number of ATAC-Seq peaks annotated to the gene. Data is shown from the wing, eye, and brain, either for all genes (open bars) or cell cycle genes (striped bars). (**E**) Line plots showing the average ATAC-Seq signal across tissues and time points at peaks associated with cell cycle genes (+/- 1 kilobase from peak center). (**F**) Summary of motif enrichment analyses on peaks associated with cell cycle genes in each tissue, including motifs annotated in insects and vertebrates. Table includes motif name and class and position weight matrix (PWM). Each motif received a Benjamini-corrected q-value < 0.05 for at least one tissue. Full motif enrichment data are available in Supplementary Table 1.

The maintenance of accessible chromatin at cell cycle genes during and after cell cycle exit suggests an active repression mechanism whereby some factor(s) continue to occupy these regions to prevent ectopic transcript expression in post-mitotic cells. To investigate what these factors might be, we performed insect and vertebrate motif enrichment analyses on the ATAC peaks nearest to cell cycle genes in each tissue (Fig. 2F and Supp. Table 1). We found that most of the significantly enriched insect motifs correspond to annotated *Drosophila* promoter sequences. This is unsurprising, given that most peaks associated with cell cycle genes are localized to promoter regions. Insect transcription factor motifs that are enriched include: M1BP, a zinc finger factor which binds the Motif 1 promoter sequence and has been implicated in transcriptional pausing, insulator functions, cellular metabolism and homeostasis (Bag et al., 2021; Chimata et al., 2023; Li and Gilmour, 2013; Poliacikova et al., 2023); DREF, a BED-finger factor that is known to regulate proliferation and other developmental processes (Killip and Grewal, 2012; Tue et al., 2017); Trl (GAGA factor), a pioneer factor thought to regulate nucleosomal as well as higher-order chromatin organization (Chetverina et al., 2021; Gaskill et al., 2021; Li et al., 2023); and an E-box motif, previously found to be upstream of *Drosophila* core promoters and similar to the binding motif for Myc, a well-described cell cycle regulator (FitzGerald et al., 2006; Qi et al., 2022). Notably, analysis of vertebrate motifs revealed enrichment for multiple annotated E2F motifs. This is again expected, given that E2F factors are evolutionarily conserved, master transcriptional regulators of the cell cycle, activating the expression of hundreds of cell cycle genes, and are known to frequently bind the promoter regions of target loci (Fischer et al., 2022; Xu et al., 2007). E2F-mediated transactivation is regulated by interactions with the retinoblastoma (Rb) protein, which binds to and represses E2Fs in a Cdk-sensitive manner. Indeed, it is thought that in postmitotic cells with low Cdk activity levels, Rb-bound E2F complexes continue to occupy binding sites and may actively repress transcript expression. Therefore, repressive Rb/E2F complexes serve as likely candidates to explain long-term chromatin accessibility after cell cycle genes have been downregulated.

The finding that accessible regions near cell cycle genes remain accessible after prolonged cell cycle exit and harbor E2F binding motifs raised the question of whether ectopic E2F activity could re-activate these transcriptional targets in postmitotic tissues after cell cycle exit has occurred. To test this, we employed the *pcna*-GFP reporter described above (Fig. 1) as a transcriptional readout of E2F activity and used the Gal4/UAS system to ectopically provide activator E2F complexes in eyes or wings, specifically after 24h APF. To ensure that this manipulation was limited to postmitotic stages, we used the “flipout” approach where a temporally controlled heat shock is used to induce expression of flippase enzyme that will catalytically remove an intervening stop codon to activate Gal4 expression. Using this approach we can limit robust heat shock-specific Gal4 activity to 40-44h APF (Fig. 3). We observed that ectopic E2F activator expression was sufficient to induce the *pcna*-GFP reporter, even in postmitotic tissues. This induction could be strengthened by the addition of ectopic Cyclin D and Cdk4, which form a G1 Cyclin/Cdk complex to inhibit Rb and further convert repressive E2F complexes to activator complexes. These data suggest that E2F-responsive regulatory elements are occupied and repressed by E2F/Rb after cell cycle exit but that they continue to be responsive to activator E2F complexes. This is consistent with an active repression model where the binding of repressor E2F/Rb complexes is maintained in postmitotic tissues to ensure transcriptional silencing even long after cell cycle exit has occurred.

**Figure 3.**
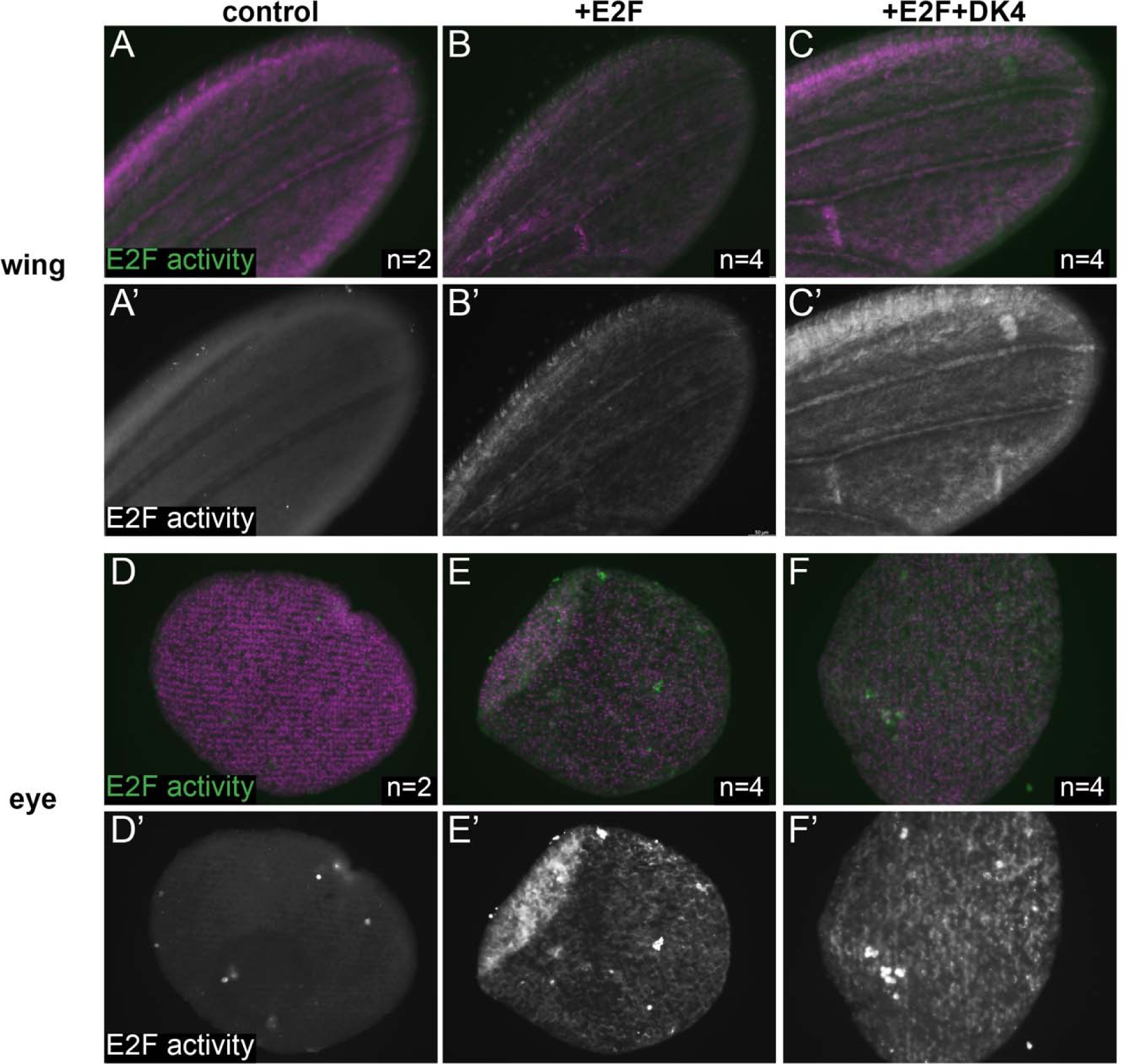
An E2F regulated accessible enhancer remains activatable after 24h APF. Wings (**A-C**) and eyes (**D-F**) were dissected from staged pupa and stained for E2F transcriptional activity (anti-GFP for PCNA-GFP reporter) at the time points indicated. Animals were collected as white pre-pupa (0h APF), staged to a postmitotic stage of 24-28h APF and heat-shocked for 20 min. to induce Gal4 expression driving E2F (UAS-E2F1+UAS-DP) or CycD+E2F (UAS-Cyclin D + UAS-Cdk4 + UAS-E2F1+UAS-DP) postmitotically. Tissues were collected and stained at 40-44h APF.

### Long-lived postmitotic fly tissues decommission enhancers at select, rate-limiting cell cycle genes

In contrast to the observations made at the majority of cell cycle genes, where accessible chromatin was limited to TSS-adjacent regions that were relatively static during and after cell cycle exit, our previous work in the wing identified three cell cycle genes that exhibited more complex regulatory architectures. The loci encoding *e2f1*, *cycE*, and *stg* - each of which act as rate-limiting components of the cell cycle - were found to harbor many candidate regulatory elements in intronic or intergenic regions (Ma et al., 2019). Many of these elements underwent apparent decommissioning (loss of accessibility) after cell cycle exit in the wing, suggesting that regulated accessibility at these critical rate-limiting genes may contribute to maintenance of the postmitotic state. To address whether these findings are generally applicable to other *Drosophila* tissues, we next compared how chromatin accessibility at these three genes changes during and after cell cycle exit across the wing, eye, and brain. At *e2f1,* peaks of accessibility span the large intronic regions of the locus (Fig. 4A). Many of these elements are shared across tissues and show similar temporal dynamics, with either maintained accessibility or apparent decommissioning by the 44h APF time point. Our findings were similar at *cycE,* where intronic elements that had been observed in the wing are also accessible in the eye and brain (Fig. 4B). Finally, the *stg* locus has previously been shown to be regulated by distal, intergenic regulatory elements (Andrade-Zapata and Baonza, 2014; Lehman et al., 1999; Lopes and Casares, 2015). Our previous work in the wing revealed that many of these regions undergo decommissioning after cell cycle exit, and we observe in our current analysis that many of the same regions show accessibility and similar dynamics in the eye and brain as well (Fig. 4C). Interestingly, in addition to accessible elements that are common across tissues, at each of these loci we are also able to discern tissue-specific dynamic elements. For example, in the larval brain we observe prominent intronic accessibility at the *cycE* locus corresponding with previously annotated enhancers that are active in the embryonic nervous system (Jones et al., 2000). These data suggest a) complex and dynamic enhancer architectures regulate the expression of a small number of rate-limiting cell cycle genes, b) the candidate regions that may be regulating these genes include both shared and tissue-specific elements, and c) enhancer decommissioning at these genes may be a common mechanism of ensuring the maintenance of cell cycle shut off across terminally differentiating tissues.

**Figure 4.**
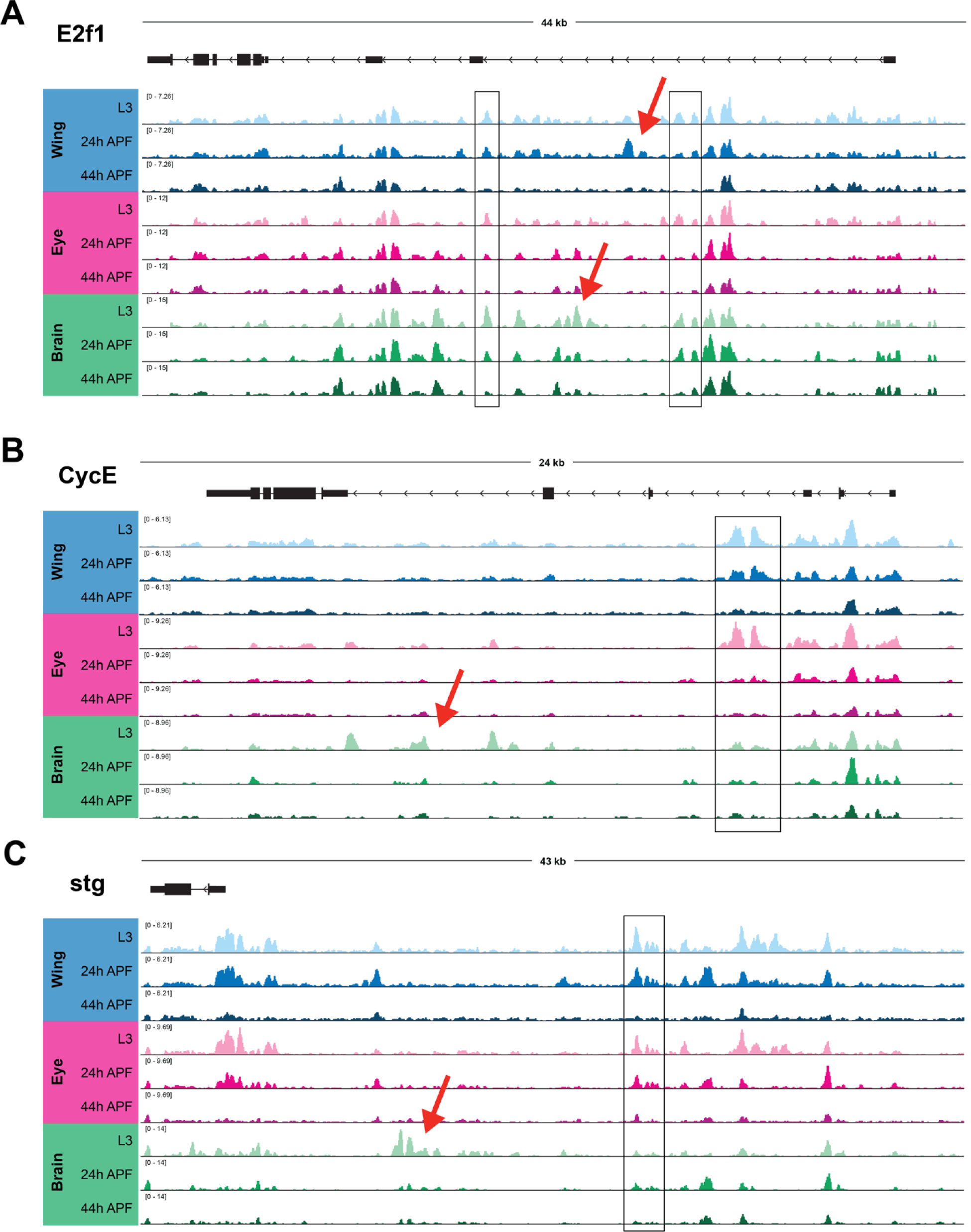
Long-lived postmitotic fly tissues decommission enhancers at select, rate-limiting cell cycle genes. ATAC-Seq accessibility data at *e2f1* (**A**), *cycE* (**B**) and *stg* (**C**) gene loci across tissues and time points. Arrows on gene diagrams indicate the direction of transcription. Y-axes indicate the normalized read counts per million. Boxes highlight example regions of shared accessibility across two or more tissues. Red arrows highlight example regions of tissue-specific accessibility.

### Cell cycle genes harbor ‘housekeeping’ and ‘developmental’ type enhancers

In genome-wide enhancer identification studies (*i.e.*, STARR-Seq), differential modes of gene regulation have been identified, employing housekeeping-type promoters and enhancers for genes that are ubiquitously expressed and developmental-type promoters and enhancers for genes that are dynamic (Zabidi et al., 2015). Given our identification of simple and complex cell cycle genes that harbor static or dynamic regulatory elements, we wondered whether analyzing STARR-Seq-defined enhancers may provide functional confirmation of enhancer activity at accessible elements as well as provide insight into the modes of regulation governing the expression of simple and complex cell cycle genes. To address this question, we generated a union set of all ATAC-Seq peaks associated with cell cycle genes in wing, eye and brain, and assessed which of those peaks overlapped with enhancers identified via STARR-Seq in ovarian somatic cells (OSC) (Zabidi et al., 2015). This revealed that 35% of cell cycle gene-associated ATAC peaks colocalize with STARR-Seq enhancers, with 20% identified as housekeeping type enhancers, 5% as developmental type enhancers, and 10% identified in both the housekeeping and the developmental datasets (Fig. 5A). It is perhaps unsurprising that most of the enhancers associated with cell cycle genes are classified as housekeeping type enhancers, as expression analyses performed in the STARR-Seq study revealed that the genes associated with housekeeping type enhancers are often widely expressed and included an enrichment of cell cycle genes (Zabidi et al., 2015). Furthermore, promoter motifs that were enriched at STARR-Seq-defined housekeeping gene TSSs were also identified in our studies as enriched within accessible regions at cell cycle genes (Fig. 2F). However, we were intrigued by the numbers of peaks at cell cycle genes co-localizing with developmental enhancers and those identified in both datasets, and wanted to further investigate the properties of these different groups of elements. It had previously been recognized that housekeeping type elements frequently localize to promoters, while developmental enhancers are enriched in intronic and intergenic regions (Zabidi et al., 2015). Therefore, we assessed the genomic distributions of cell cycle-associated peaks of each type. As described above, ATAC-Seq peaks associated with cell cycle genes are enriched for promoter regions relative to all genome-wide peaks (Fig. 2C), with approximately 60% at promoters, 20% in introns, and small numbers mapping to exons, downstream, and intergenic regions. Consistent with previous findings, we found that peaks overlapping with housekeeping and/or developmental STARR-Seq enhancers show differential distributions, with almost all of the peaks overlapping housekeeping elements localizing to promoters and those overlapping developmental enhancers showing a relative depletion for promoter regions and enrichments for intronic regions and regions just downstream of gene bodies (Fig. 5B). Moreover, the ATAC-Seq peaks that showed overlap with both housekeeping and developmental type enhancers show a distribution reminiscent of housekeeping elements, where almost all of these peaks localize to promoter regions.

**Figure 5.**
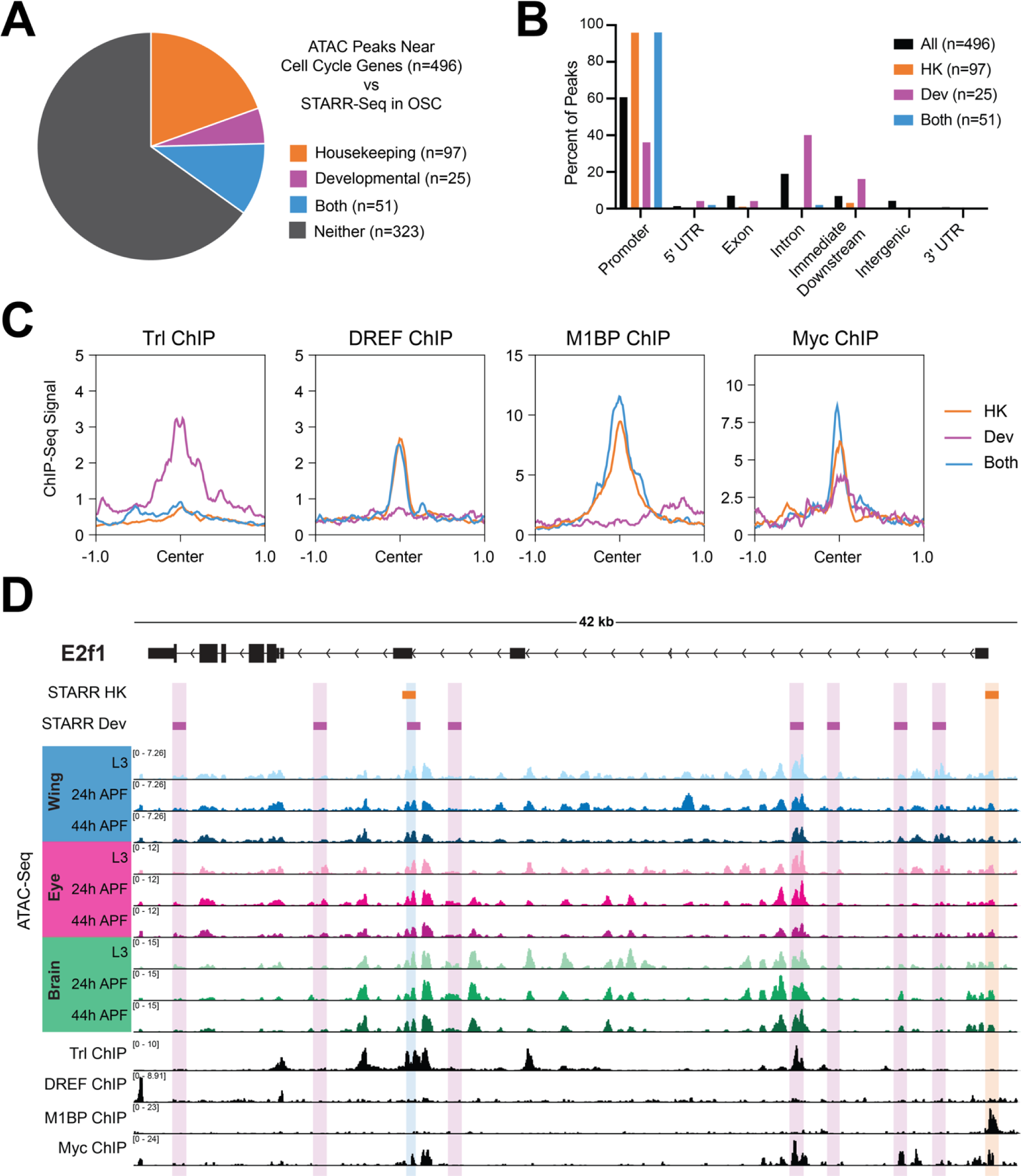
Cell cycle genes harbor ‘housekeeping’ and ‘developmental’ type enhancers. (A) Pie chart depicting overlap between the union set of ATAC-Seq peaks associated with cell cycle genes across the wing, eye, and brain (n=496) and STARR-Seq identified enhancers from ovarian somatic cells (OSC). (B) Bar chart showing the genomic distributions of ATAC-Seq peaks at cell cycle genes: for all peaks (black), those overlapping with housekeeping enhancers (orange), overlapping with developmental enhancers (magenta), or overlapping with both datasets (blue). HK, housekeeping. Dev, developmental. (C) Line plots showing the average ChIP-Seq signal for Trl, DREF, M1BP, and Myc at cell cycle gene-associated ATAC-Seq peaks (+/- 1 kilobase from peak center) for those overlapping with housekeeping enhancers (orange), overlapping with developmental enhancers (magenta), or overlapping with both datasets (blue). HK, housekeeping. Dev, developmental. (D) STARR-Seq, ATAC-Seq, and ChIP-Seq data at the *e2f1* locus. Orange boxes indicate STARR-Seq housekeeping enhancers and magenta boxes indicate STARR-Seq developmental enhancers. ATAC-Seq accessibility data from wing, eye, and brain at L3, 24h APF, and 44h APF. ChIP-Seq data for Trl, DREF, M1BP, and Myc. Y-axes indicate the normalized read counts per million. HK, housekeeping. Dev, developmental.

Besides the differences in genomic distribution, STARR-Seq enhancers also show differences in motif composition. Housekeeping enhancers are enriched for DREF motifs, while developmental elements show a broader diversity of enriched motifs, including Bap, Apterous, and Trl (GAGA factor) among others (Zabidi et al., 2015). Importantly, there appears to be a functionally important distinction between DREF and Trl binding in the determination of housekeeping versus developmental enhancers; it was shown that artificially exchanging these motifs was sufficient to change the profile of housekeeping versus developmental type activity in individual enhancers (Zabidi et al., 2015). Both of these motifs had been identified in our analysis of enriched motifs among cell cycle-associated peaks (Fig. 2F). Therefore, we wondered whether binding of DREF and Trl may also functionally differentiate elements associated with cell cycle genes. To test this, we analyzed Trl ChIP-Seq data generated in larval wing discs (Oh et al., 2013) and DREF ChIP-Seq data generated in Kc cells (Gurudatta et al., 2013), and assayed the binding of these factors at ATAC peaks associated with housekeeping, developmental or both types of enhancers. Consistent with previous findings, we found that Trl signal is enriched at developmental type elements and DREF preferentially binds at housekeeping type elements (Fig. 5C). Interestingly, as with the genomic distributions, peaks associated with both classes are more reminiscent of housekeeping elements: these are not generally bound by Trl and are bound by DREF. In addition to DREF and Trl, motifs for other factors enriched among cell cycle-associated peaks in our dataset included M1BP and an E-box motif, possibly regulated by Myc (Fig. 2F). We wondered whether these factors also show preferential binding at housekeeping or developmental enhancers, so we performed the same analysis using M1BP (Bag et al., 2021) and Myc (Yang et al., 2013) ChIP-Seq data generated in Kc cells. This revealed that M1BP shows preferential binding at peaks corresponding to housekeeping elements and those identified as active in both datasets, and does not bind strongly at peaks associated with developmental enhancers (Fig. 5C). This is not surprising, given that M1BP binds to the Motif 1 promoter sequence and shows the strongest binding at the classes of elements localizing to promoters. Finally, Myc ChIP-Seq showed that Myc binds to some degree at each group of peaks (Fig. 5C), suggesting that Myc may act more broadly across both types of enhancers. As a whole, these analyses support the previous assertion than many elements associated with cell cycle genes show housekeeping type activity, but clarify that cell cycle genes also harbor elements with developmental type activity as well as elements of both sets, suggesting that there may be more nuance to cell cycle gene regulation than strictly ubiquitous, housekeeping-type regulation. This is unsurprising, given that cell cycle genes are expressed in a developmentally dynamic manner (Fig. 2B) and must be repressed at the proper time for cell cycle exit and proper development to proceed (Du et al., 1996; Ma et al., 2019; Pilaz et al., 2009; Ruijtenberg and van den Heuvel, 2016; Tarui et al., 2005).

Finally, we were particularly interested to assess the presence of STARR-Seq enhancers identified at the three rate-limiting cell cycle genes that we had identified as having complex, dynamic regulatory landscapes: *e2f1*, *cycE*, and *stg*. We found that a number of STARR-Seq enhancers map to the *e2f1* locus, including housekeeping and developmental enhancers, as well as one element identified in both datasets (Fig. 5D). Some of these elements overlap accessibility peaks identified by ATAC-Seq in the wing, eye, and brain, and some of these elements overlap with ChIP-Seq data for Trl, DREF, M1BP, and/or Myc. However, we noted that there is not a great deal of overlap between enhancers identified by STARR-Seq (in cell culture) and the ATAC-Seq peaks that are most developmentally dynamic or tissue specific *in vivo*. Similar findings were made at the *cycE* and *stg* loci and were supported by an analysis of developmentally dynamic versus static peaks genome-wide (Supp. Fig. 1). These data suggest that developmental factors and/or processes driving gene regulatory events *in vivo* are not well recapitulated in the cell culture models that are required for high-throughput, genome-wide enhancer identification, and support a need for studies of candidate elements *in vivo* to understand the regulation of developmentally dynamic genes.

### Dynamic chromatin regions within *e2f1* and *stg* show enhancer activity

To confirm which dynamically accessible regions at the complex cell cycle gene loci serve as bona fide *in vivo* enhancers during metamorphosis, we searched the publicly available Janelia Flylight and Vienna Tile collections for Gal4 driver lines that overlap with dynamically accessible regions at these loci (Kvon et al., 2014; Pfeiffer et al., 2008). Although there were no Gal4 driver lines that would provide information about the dynamic regions at *cycE*, we note that some of the dynamic regions that we identified at this locus have previously been validated as enhancers in other tissues and/or time points (Deb et al., 2008; Djiane et al., 2013; Jones et al., 2000; Kannan et al., 2010). We selected five driver lines of interest at *e2f1* (Fig. 6A) and seven lines of interest at *stg* (Fig. 7A). To visualize the activity of these drivers, we crossed each of them to G-TRACE (Evans et al., 2009), allowing us to assess current Gal4 expression using UAS-RFP as well as past driver activity by permanent GFP expression in cells that expressed Gal4 at any point in their developmental lineage. Most of these driver lines were assessed in both the eye and wing at L3, 24h APF, and 44h APF time points; many lines were also tested in the brain. Of the five drivers tested from *e2f1*, four show enhancer activity in at least one tissue. Similarly, of the seven drivers tested from the *stg* regulatory region, all show enhancer activity in at least one tissue. Note that we selected a subset of driver lines to highlight in Figures 6B-D and 7B-D, but comprehensive data for all drivers in all tissues tested are included in Supplementary Figures 2-7.

**Figure 6.**
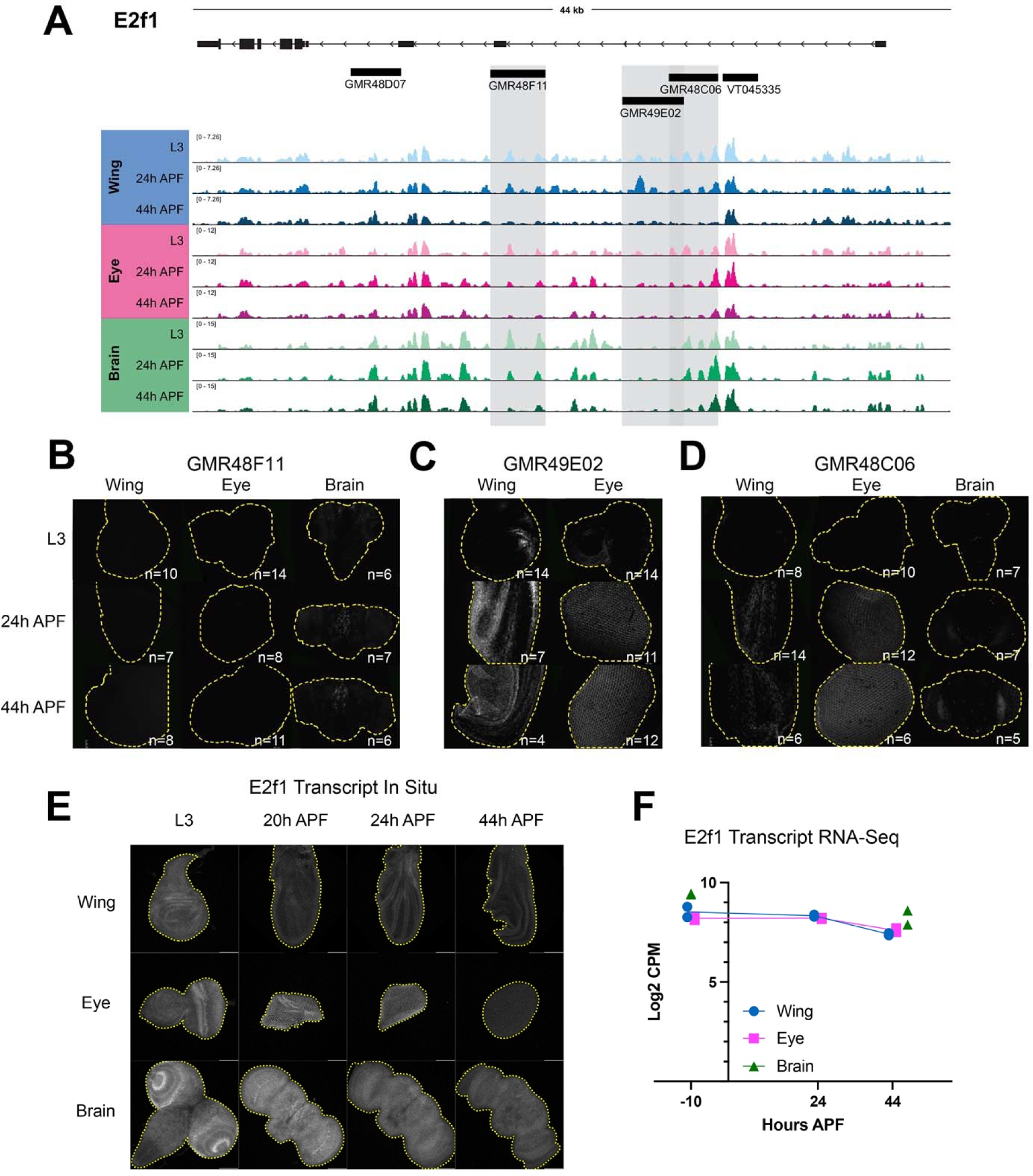
Dynamic chromatin regions within *e2f1* show enhancer activity *in vivo*. (**A**) ATAC-seq accessibility data at the *e2f1* locus across tissues and timepoints. Fragments used to direct Gal4 expression in publicly available driver lines are depicted by black bars, each of which was tested for enhancer activity. (**B-D**) Enhancer expression data are presented for driver lines GMR48F11, GMR49E02, and GMR48C06, which are highlighted by gray boxes in (A). Each driver was tested in wing, eye and/or brain at L3, 24h APF and 44h APF time points. Images show the readout of ‘current’ Gal4 activity (UAS-RFP) and signal intensities are qualitative to emphasize the distinct spatial domains of activity across driver lines. Comprehensive data from all drivers is available in Supp. Figs 2-4. (**E**) HCR-FISH data showing *e2f1* transcript expression in wing, eye and brain at L3, 20h, 24h, and 44h APF. (**F**) Line plot showing e2f1 Log2-transformed counts per million (CPM) expression levels via RNA-Seq in wing and eye at L3, 24h and 44h APF timepoints, and in brain at L3 and 48h APF timepoints. Individual data points represent values from RNA-Seq replicates.

**Figure 7.**
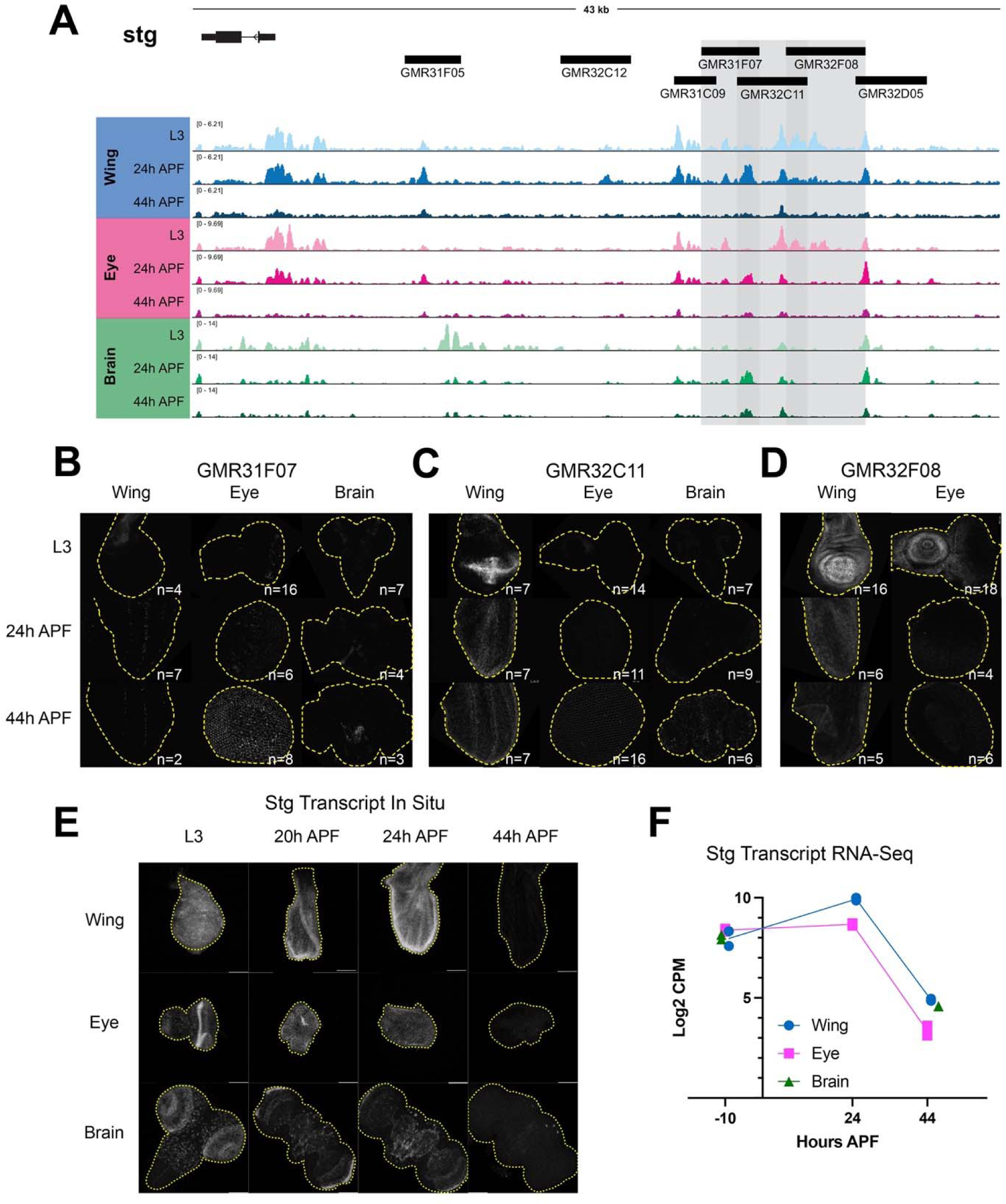
Dynamic chromatin regions in the *stg cis-*regulatory region have enhancer activity *in vivo*. (**A**) ATAC-seq accessibility data at the *stg* locus across tissues and timepoints. Fragments used to direct Gal4 expression in publicly available driver lines are depicted by black bars, each of which was tested for enhancer activity. (**B-D**) Enhancer reporter expression data are presented for driver lines GMR31F07, GMR32C11, and GMR32F08, which are highlighted by gray boxes in (A). Each driver was tested in wing, eye and/or brain at L3, 24h APF and 44h APF time points. Images show the readout of ‘current’ Gal4 activity (UAS-RFP) and signal intensities are qualitative to emphasize the distinct spatial domains of activity across driver lines. Comprehensive data from all drivers is available in Supp. Figs. 5-7. (**E**) HCR-FISH data showing stg transcript expression in wing, eye and brain at L3, 20h, 24h, and 44h APF. (**F**) Line plot showing stg Log2-transformed counts per million (CPM) expression levels via RNA-Seq in wing and eye at L3, 24h and 44h APF timepoints, and in brain at L3 and 48h APF timepoints. Individual data points represent values from RNA-Seq replicates.

Overall, our enhancer studies at *e2f1* and *stg* led us to a number of important observations regarding the principles of regulation at these complex cell cycle genes. First, some drivers show enhancer activity in multiple tissues, while a number of driver regions show tissue-specific enhancer activity. Examples of tissue-specific elements include GMR48F11 at *e2f1*, which is active in the brain (Fig. 6B), and the *stg* driver GMR32C12 which shows activity specifically in the lamina, the tissue connecting the retina to the optic lobe (Supp Fig. 6). These results highlight that even when a genomic region is accessible across tissues, enhancer activity may be regulated by factors differentially expressed in those contexts. Next, we noted that many drivers show enhancer activity in specific domains of a tissue; this is especially evident in the wing and the brain. Examples of this include GMR48C06 at *e2f1* (Fig. 6D) and GMR31F07 at *stg* (Fig. 7B). To complement the reporter expression data showing distinct spatial domains of enhancer activity, we assayed the transcript expression of e2f1 and stg in each tissue via HCR-FISH (Fig. 6E and 7E). These data are consistent with the expected temporal expression dynamics that we observe for these genes via RNA-Seq (Fig. 6F and 7F), and confirm that each transcript is widely expressed within each tissue. Taken together, these data suggest that *e2f1* and *stg* are regulated by enhancers that act in a modular manner, whereby individual enhancers activate transcription in distinct compartments that ultimately, in combination, drive widespread transcript expression across the tissue. Finally, we noted that some enhancers show the expected temporal dynamics of activity based on accessibility data, while others do not. In particular, the eye often exhibited enhancer activity at the 44h APF time point, that would not be expected based on decreasing accessibility at the corresponding driver regions. However, there are a number of possible explanations for these discrepancies. First, the resolution for temporal dynamics of enhancer reporter assays are limited by the long half-lives of standard fluorescent reporter proteins, as is the case in these experiments using RFP to read out Gal4 expression. This is supported by the more rapid dynamics of silencing observed in the wing using a de-stabilized GFP (UAS-dsGFP, Supp. Fig. 10). Second, we note some reporter expression in the late pupal eye using Janelia’s empty Gal4 control line, suggesting that there may be some factor(s) expressed in the pupa eye that induce expression of Gal4 off of the synthetic core promoter used in the generation of these drivers (Supp. Fig. 11). Finally, it is also possible that these enhancer regions do not fully recapitulate their endogenous activities and dynamics when removed from the native genomic context. Despite these limitations, we found that the intensity of reporter expression for some elements in the wing that we assessed across a finer time course do show a peak of activity around 24h APF followed by decreasing reporter intensity up to 44h APF (Supp. Fig. 8,9). This is consistent with the decommissioning or loss of accessibility that these elements exhibit via ATAC-Seq, and suggests that enhancer decommissioning may be one mechanism used to ensure cells maintain a non-cycling, postmitotic state after cell cycle exit.

### Stable repression together with decommissioning of enhancers at rate-limiting cell cycle genes ensures robust cell cycle exit

Our model for genomic control of cell cycle gene expression in *Drosophila* is as follows (Fig. 8): Cell cycle genes with simple enhancer architecture contain promoter proximal enhancers enriched for “housekeeping-associated” motifs such as E2F binding sites and DREF core promoters. These elements exhibit accessibility during proliferation as well as after cell cycle exit. The post-mitotic transcriptional repression of these genes is controlled through the E2F complex switching from an activating to repressive form, influenced by cyclin/cdk activity and the phosphorylation of Rbf. By contrast, rate-limiting cell cycle genes with complex, modular enhancer architecture such as *cyclin E*, *E2f1* or *string*, may be influenced by E2F complexes, but are also controlled via E2F-independent mechanisms through enhancers that bind other transcription factors such as Su(H) or bHLH proteins (Andrade-Zapata and Baonza, 2014; Djiane et al., 2013). These genes exhibit enhancer decommissioning after the transition to a postmitotic state to ensure their silencing despite the re-use of signaling pathways such as Notch or EGF in post-mitotic terminal differentiation processes.

**Figure 8:**
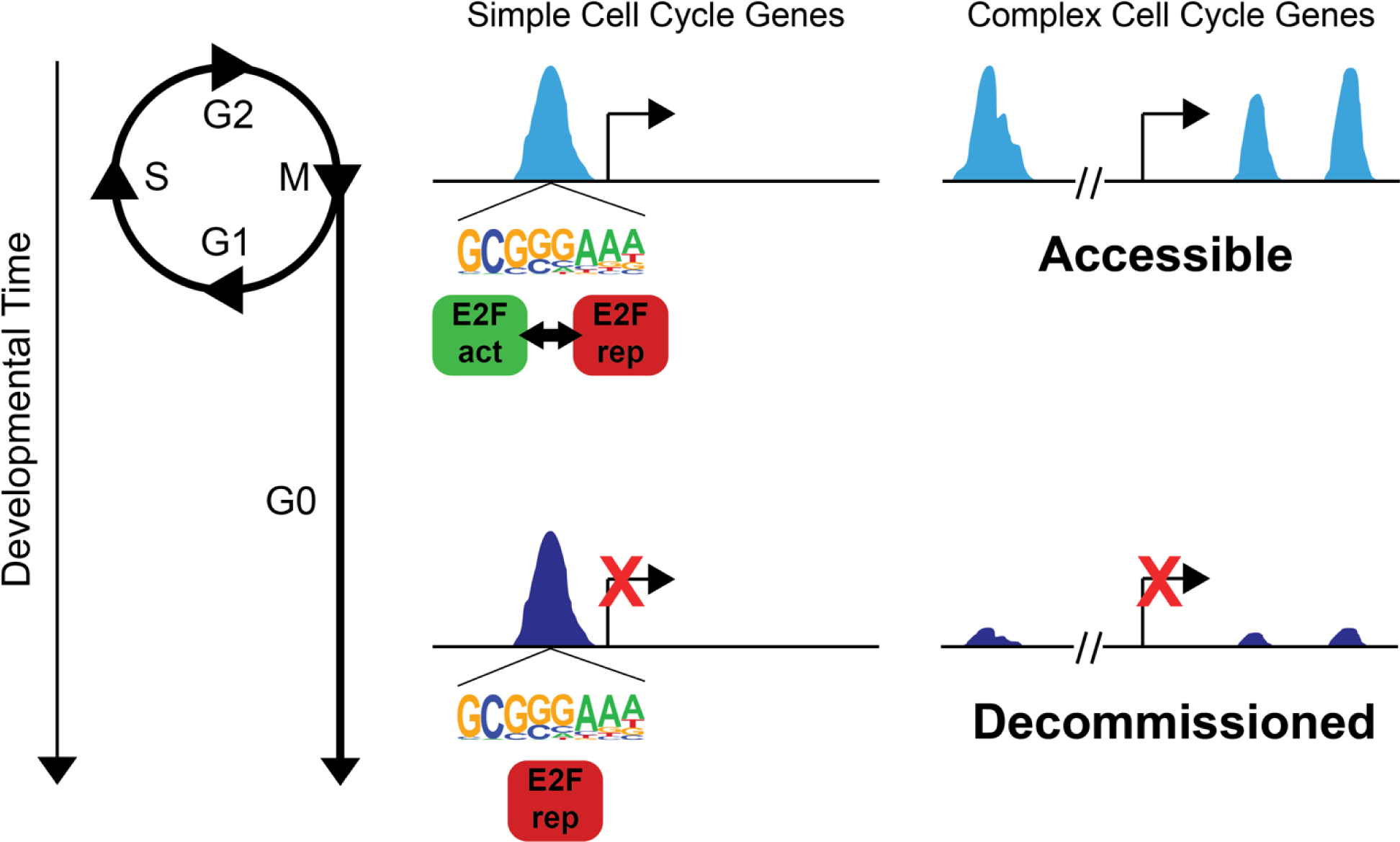
Model. Stable repression at most cell cycle genes together with decommissioning of enhancers at rate-limiting cell cycle genes ensures robust cell cycle exit. Cell cycle genes with simple enhancer architecture contain promoter proximal enhancers enriched for E2F binding sites that exhibit accessibility during proliferation as well as after cell cycle exit. These sites are dynamically occupied by activator or repressor complexes in a cell cycle phase-dependent manner in proliferative cells. After cell cycle exit, they are stably occupied by repressor complexes to maintain long-term repression. Rate-limiting cell cycle genes with complex, modular enhancer architecture exhibit enhancer decommissioning after the transition to a postmitotic state.

### Principles of Cell Cycle Gene Regulation are Conserved in Mammalian Retina

We next wondered whether the model that we propose for cell cycle gene regulation in *Drosophila* is evolutionarily conserved and applicable to mammalian tissues. To test this, we examined a published RNA-Seq and ATAC-Seq dataset from the mouse retina spanning the developmental trajectory of this tissue from proliferation to cell cycle exit and terminal differentiation (Aldiri et al., 2017). New cells are born in the mouse retina at the highest rates between postnatal (P) days P0 and P3, after which the proliferative rate decreases with the tissue becoming fully postmitotic by P10 (Aldiri et al., 2017; Bremner et al., 2004). By P21, the retina has undergone terminal differentiation, generating seven classes of mature retinal cells. First, we assessed cell cycle gene expression across this time course via RNA-Seq. Taking the list of cell cycle genes in *Drosophila* (Fig. 2B) and collecting all of the orthologs of those genes, we recovered 564 mouse cell cycle genes that were expressed in at least one sample of the postnatal retina time course. We found that many of these genes show variable temporal dynamics, while many others exhibit the expected pattern of core cell cycle components: high expression early that decreases during and after cell cycle exit (Fig. 9A). In particular, there are 228 cell cycle genes that exhibit a continual decrease in transcript expression level between each sequential time point, which we refer to as ‘repressed’ cell cycle genes.

**Figure 9:**
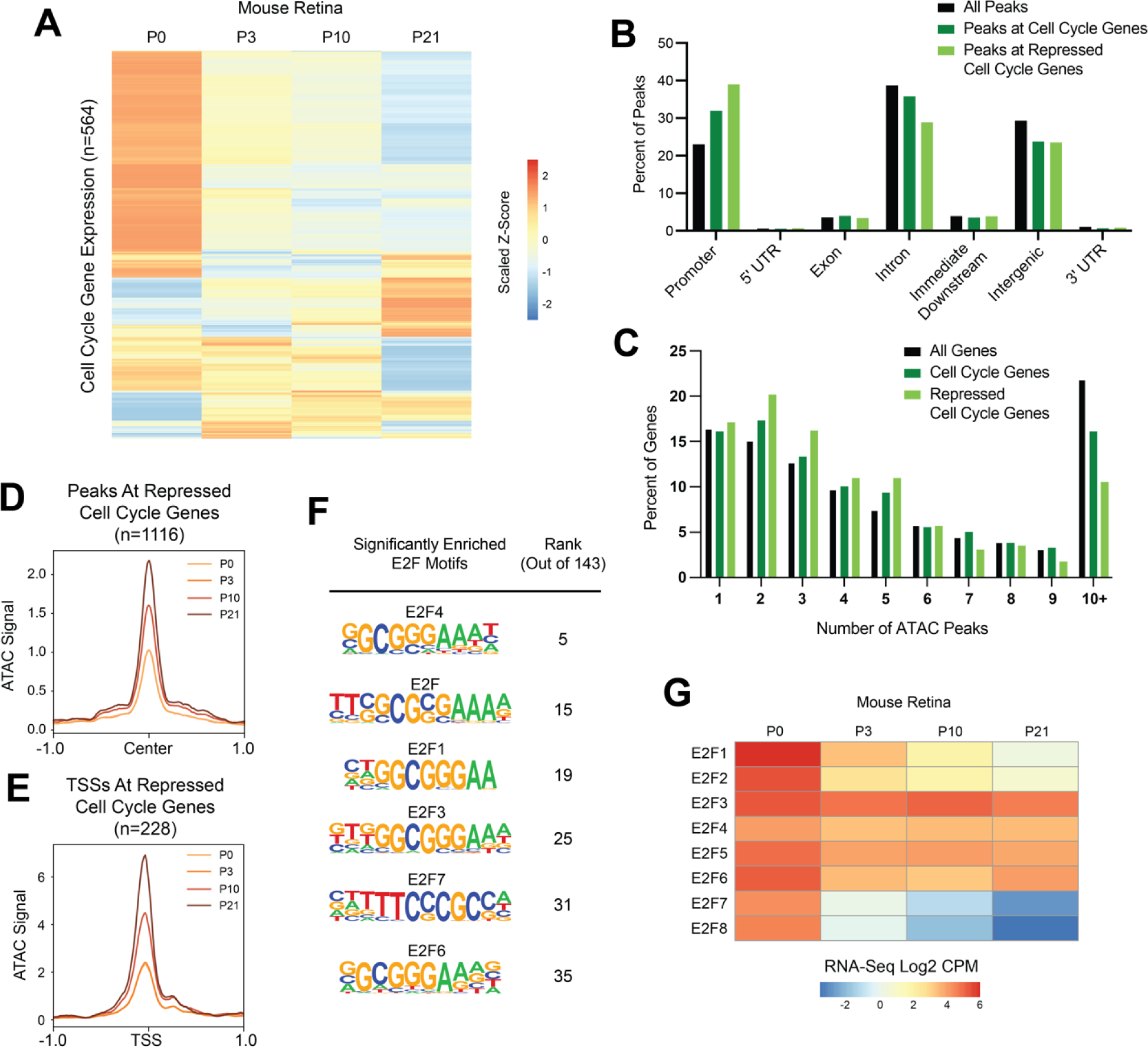
Many cell cycle genes in mouse retina have simpler than average regulatory architecture, are transcriptionally repressed after cell cycle exit, but retain chromatin accessibility during terminal differentiation. (**A**) Heatmap depicting the average transcript expression values for 564 genes with annotated functions related to the cell cycle. Data are scaled by Z-score and hierarchically clustered. (**B**) Bar plot showing the genomic distributions of ATAC-Seq peaks from the mouse retina, either for all peaks (black bars), peaks associated with cell cycle genes (dark green bars), or peaks associated with repressed cell cycle genes (light green bars) (**C**) Bar plot showing the distributions of genes binned by number of ATAC-Seq peaks annotated to the gene. Data is shown from the mouse retina for all genes (black bars), cell cycle genes (dark green bars), or repressed cell cycle genes (light green). (**D**) Line plot showing average ATAC-Seq signal at peaks (+/- 1 kilobase from peak center) associated with the repressed cell cycle genes. P0, light orange. P3, orange. P10, dark orange. P21, brown. (**E**) Line plot showing average ATAC-Seq signal at TSSs (+/- 1 kilobase) for continually decreasing cell cycle genes. P0, light orange. P3, orange. P10, dark orange. P21, brown. (**F**) Enrichment of E2F motifs in peaks associated with repressed cell cycle genes. Table includes motif name and position weight matrix (PWM) and rank among the 143 motifs that were significantly enriched. (**G**) Heatmap depicting the average transcript expression values for E2F family genes. Data are presented as normalized Log2 Count Per Million (CPM) values.

The first part of our model states that most cell cycle genes exhibit a simple regulatory architecture and are regulated primarily by promoter-proximal elements. These elements retain accessibility after cell cycle exit and transcriptional repression, and are enriched for E2F and ‘housekeeping’ motif sequences. To investigate this component of the model in mouse retina, we first compared the genomic distributions of ATAC-Seq peaks genome-wide, peaks associated with cell cycle genes, and peaks associated with repressed cell cycle genes. This revealed that as in Drosophila, mouse cell cycle genes show an enrichment of accessible chromatin localizing to promoters and a smaller proportion of peaks localizing to intronic and intergenic regions relative to the genome-wide distribution (Fig. 9B). Promoter enrichment and intronic depletion were even more pronounced among peaks associated with repressed cell cycle genes. Next, we analyzed the number of ATAC-Seq peaks assigned per gene as a measure of the complexity of the regulatory architecture. This revealed that as in *Drosophila*, mouse cell cycle genes exhibit a simpler than average architecture that is most pronounced among the repressed genes, with an enrichment of genes harboring just two or three accessible regions, as well as a depletion of very complex loci harboring 10 or more ATAC-Seq peaks (Fig. 9C). To address the component of the model that argues for maintained chromatin accessibility in the post-mitotic state, we assessed the time course of ATAC-Seq data at peaks associated with repressed cell cycle genes as well as at the TSSs of those genes. We chose to analyze ATAC-Seq data associated with only the repressed cell cycle genes to exclude the possibility that the accessibility profile would be influenced by genes whose expression was increasing, fluctuating, or remaining constant over the time course. This revealed that on average, peaks and TSSs associated with repressed cell cycle genes increased in accessibility at the P10 and P21 postmitotic time points (Fig. 9D-E). However, this increase in accessibility was seen across all peaks genome-wide at the later time points (data not shown), leading us to conclude that this finding does not represent a regulatory process that is specific to cell cycle genes. Nonetheless, the maintenance of chromatin accessibility at cell cycle genes that undergo transcriptional silencing during cell cycle exit in the mouse retina supports the idea that this is a conserved feature of cell cycle gene regulation. As in *Drosophila*, we interpret this as evidence that these sites continue to be occupied in the post-mitotic state, perhaps to maintain long term transcriptional repression. To investigate what factor(s) may be present, we performed motif enrichment analysis on the peaks associated with repressed cell cycle genes, and found that there were 143 motifs significantly enriched in these regions (Supp. Table 2). These include binding sequences for factors that have been previously described to bind housekeeping type promoters in mammals, including members of the ATF, CREB, Myc, NRF, SP, and USF families (Farre et al., 2007). We were particularly interested in the enrichment of E2F family motifs, which were among the most significantly enriched (Fig. 9F). In mammals there are 8 E2F family members, some of which function as activators, others as repressors, and some with atypical properties (Fischer et al., 2022). It has previously been shown that differential expression of E2F family members is associated with proliferation versus cell cycle exit decisions (Cuitino et al., 2019). Therefore, we analyzed the transcript expression levels of all E2F family members in the mouse retina across this time course. We found that activator family members E2F1 and E2F2, as well as the atypical family members E2F7 and E2F8, are highly expressed at P0 and then are transcriptionally repressed, while others including the repressors E2F4, 5, and 6 continue to be well expressed post-mitotically (Fig. 9F). The enrichment of E2F binding motifs among candidate regulatory regions that remain accessible after cell cycle exit, along with the transcriptional profile wherein repressor E2Fs continue to be expressed after cell cycle exit, support the idea that binding of E2F repressor complexes contributes to post-mitotic transcriptional repression of target genes. As a whole, these data are consistent with the model for transcriptional regulation of simple cell cycle genes in *Drosophila,* suggesting that this is evolutionarily conserved and applicable to mammalian tissue.

We next sought to investigate whether some mammalian cell cycle genes harbor dynamic regulatory elements in the manner of *e2f1*, *stg*, and *cycE* in *Drosophila*. Given the lack of functionally validated distal regulatory elements that have been described for mammalian cell cycle genes, we expected that this question would be difficult to answer with certainty. Nonetheless, we began by analyzing ATAC-Seq data from the mouse retina at the orthologs of *e2f1*, *stg*, and *cycE.* This revealed no prominent, dynamically accessible regions at the orthologs of stg and cycE (Supp. Fig. 12), but did identify two peaks residing less than 10 kilobases upstream of *E2f1* that exhibit apparent decommissioning upon cell cycle exit (Fig. 10A-B). These elements are marked by H3K27 acetylation - a histone modification that labels active regulatory elements - in the mouse retina at the proliferative time points and this mark is lost coincident with cell cycle exit. Therefore, although these putative enhancers have not been functionally validated to regulate *E2f1*, the available data are suggestive of post-mitotic decommissioning at the *E2f1* locus in mammals and support further investigation. We next sought to leverage previous work describing the cis-regulatory landscapes of Myc family members in cancer (Helmsauer et al., 2020; Lancho and Herranz, 2018; Schuijers et al., 2018; Zaytseva and Quinn, 2017), as the transcriptional regulation of these loci are among the most well-studied among mammalian cell cycle regulators. Most notably, the *Mycn* locus has been implicated in developmental growth of the retina, where it is required for proper retinal size and coordination of retina to eye size (Martins et al., 2008), as well as the development of retinoblastomas (Lee et al., 1984; Rushlow et al., 2013). We found that *Mycn* transcripts are well expressed in the early postnatal retina and strongly downregulated by P21 (Fig. 10C). This transcriptional downregulation is accompanied by loss of chromatin accessibility and H3K27 acetylation across the gene body after cell cycle exit. Interestingly, a number of far distal putative enhancers have been implicated in Mycn expression in the context of neuroblastoma (Helmsauer et al., 2020). We therefore expanded our analysis to the intergenic regions surrounding this locus and found further evidence of candidate regulatory elements that lose accessibility at the post-mitotic time points, some of them corresponding to elements identified in neuroblastoma cells (Fig. 10D). A number of these regions exhibit loss of H3K27ac after cell cycle exit, while a large region, hundreds of kilobases from the Mycn locus is targeted for repressive H3K27me3 deposition at the postmitotic timepoints. Similar findings were made at other Myc family member genes (Supp. Fig. 13). While these findings require functional validation of the putative enhancer elements, they are again strongly suggestive of decommissioning at the *Mycn* locus in the mouse retina after cell cycle exit.

**Figure 10:**
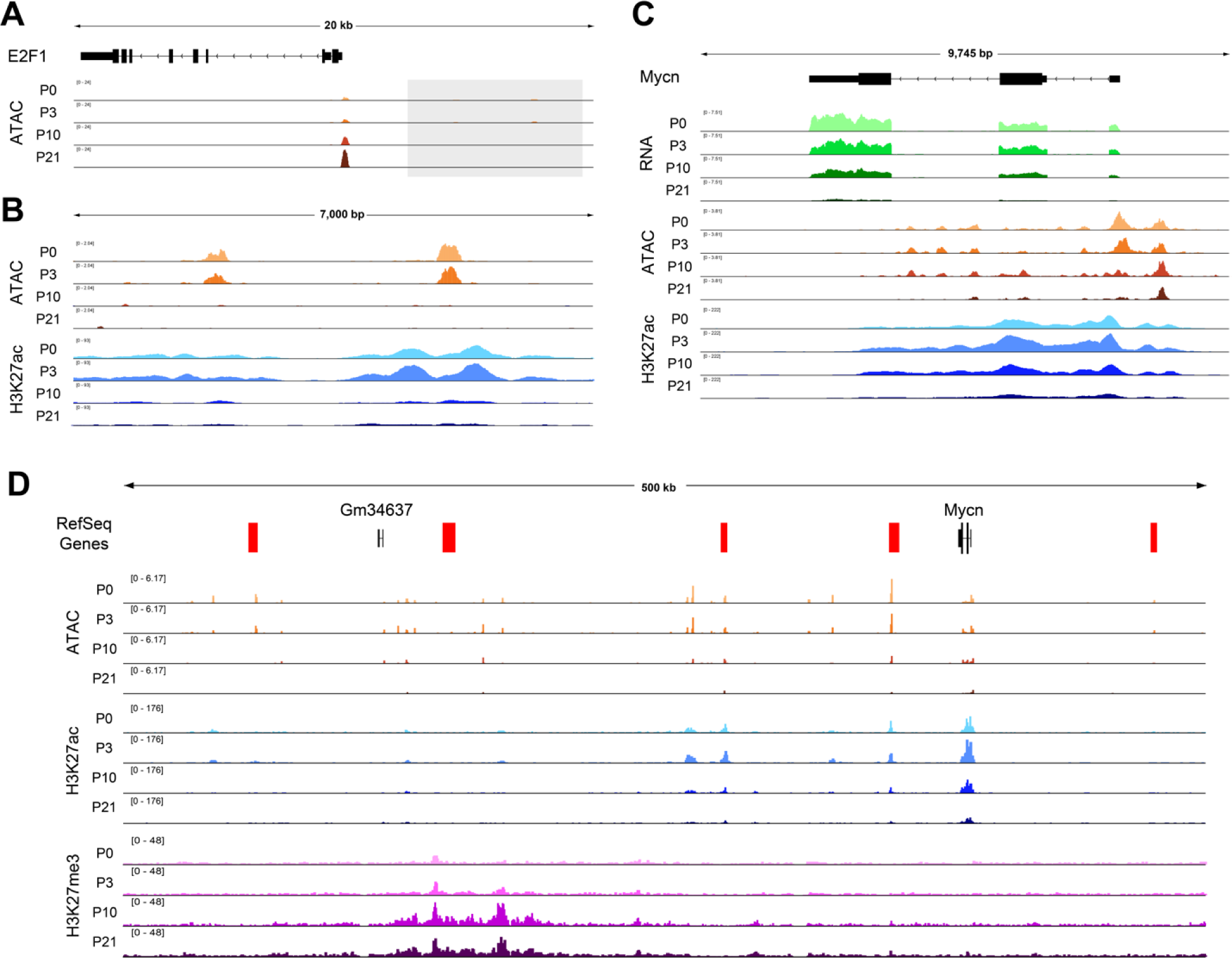
Putative enhancers at E2F1 and MycN suggest postmitotic decommissioning at some cell cycle genes in mouse retina. (**A**) ATAC-Seq data tracks from the mouse retina at the *E2f1* locus. The gray box indicates the upstream region that is depicted in panel B. (**B**) The intergenic region upstream of *E2f1*, showing tracks for ATAC-Seq and H3K27ac ChIP-Seq data from mouse retina. (**C**) The *Mycn* gene body, with RNA-Seq, ATAC-Seq, and H3K27ac ChIP-Seq data from mouse retina. (**D**) A view of the large intergenic regions surrounding the *Mycn* locus, with ATAC-Seq, H3K27ac ChIP-Seq, and H3K27me3 ChIP-Seq data from mouse retina.

In sum, the data at *E2f1* and *Mycn* are supportive of a conserved cell cycle control mechanism that is shared between *Drosophila* and mammals, whereby genes encoding critical cell cycle regulatory components undergo decommissioning to prevent spurious transcript expression in the post-mitotic state.

## Methods

### E2F transcriptional reporter assays

The PCNA-GFP reporter with characterized E2F binding sites is described in (Thacker et al., 2003). For Figure 1, genotypes were: *w; + ; PCNA-GFP* with animals aged at 25°C. For Figure 3 genotypes were:

Control: *y,w,hs-flp/w; UAS-RFP/+; PCNA-GFP, act>stop>Gal4* +E2F: *y,w,hs-flp/w; UAS-RFP/UAS-E2F1, UAS-Dp; PCNA-GFP, act>stop>Gal4/+* E2F+DK4: *y,w,hs-flp/w; UAS-RFP/UAS-CycD, UAS-Cdk4; PCNA-GFP, act>stop>Gal4/ UAS-E2F1, UAS-Dp* UAS-E2F and UAS-DP are from BDSC 4774 UAS-CycD,UAS-Cdk4 are from (Datar et al., 2000) y,w,hsflp is from BDSC 1929 act>stop>Gal4 on III is from: BDSC 4780

Animals were heat-shocked for 20 min at 37°C at 24-28h APF and dissected at 40-44h APF.

### Immunofluorescence

Tissues were fixed in 4% Paraformaldehyde/PBS and stained as described (Ma et al., 2019). Mitoses were assayed using rabbit anti phospho-histone H3 (PH3), (Cell Signaling) at 1:1000. Anti-GFP staining was performed with rabbit anti-GFP antibody (Invitrogen) at 1:1000.

Secondaries were Alexa-488 or Alexa564 conjugated goat anti-rabbit (Invitrogen) used at 1:2000. DNA was stained with Dapi (Sigma). Tissues were mounted on slides with Vectashield mounting medium and imaged with a Leica SP5 confocal microscope.

### ATAC-Seq and RNA-Seq Sample and Library Preparation

Genotypes and staging: Wings, eyes, and brains were dissected from w1118/y,w,hsflp122 ; +; + female animals for all samples. Animals were raised at room temperature on Bloomington Cornmeal media without malt extract (bdsc.indiana.edu/information/recipes/bloomfood.html). Larval samples were dissected from wandering larvae isolated from uncrowded vials. Vials with more than ∼100 larvae were diluted into fresh vials to keep larvae uncrowded. Pupa were collected from vials at the White Pre-pupa stage (WPP) as described (Flegel et al., 2013), which was taken as 0h After Puparium Formation (APF) and incubated on damp Kimwipes at 25°C to the indicated hours APF.

ATAC-seq: Wing ATAC-Seq data were previously published and re-analyzed for the current study (Buchert et al., 2023). All samples were dissociated using collagenase/dispase prior to a standard Omni-ATAC protocol, as previously described (Buchert et al., 2023; Corces et al., 2017). 10 wings, 16 eyes, or 3 brains were used per sample. Larval eye discs were separated from antenna discs during dissection. Library quality was assessed with an Agilent Bioanalyzer or Tape Station. Wing and eye ATAC-Seq libraries were sequenced on an Illumina NovaSeq SP 100 cycle flow cell for 50 bp paired end reads, at a target depth of 90 million reads per sample. Brain ATAC-Seq libraries were sequenced on an Illumina NovaSeq S4 300 cycle flow cell for 150 bp paired end reads, at a target depth of 70 million reads per sample.

RNA-seq: Wing RNA-Seq data were previously published and re-analyzed for the current study (Ma et al., 2019). 16 eyes were used per sample. Larval eye discs were separated from antenna discs during dissection. RNA was extracted using a standard Trizol/chloroform protocol, precipitated overnight at −20C in isopropanol, and quality was assessed with Agilent Bioanalyzer. Libraries were prepared using polyA selection and again assessed with Agilent Bioanalyzer or Tape Station prior to sequencing. Libraries were sequenced on an Illumina NovaSeq SP 100 cycle flow cell for 50 bp paired end reads, at a target depth of 90 million reads per sample.

### ATAC-Seq Data Analysis

Adaptors and low-quality bases were trimmed using cutadapt 1.18 (Martin, 2011). Reads were aligned to DM6 or MM10 using Bowtie2.4.1 (Langmead and Salzberg, 2012) using --local -- very-sensitive parameters and max fragment size set to 1000bp. PCR duplicates were marked using picard-tools 2.8.1 MarkDuplicates (“Picard Toolkit.” 2019. Broad Institute, GitHub Repository. https://broadinstitute.github.io/picard/; Broad Institute). Downsampling was done to normalize read depth across samples and reads spanning less than 120 bp (sub-nucleosomal fragments) were used for analysis. BAM files were generated using samtools 1.5 (Li et al., 2009), and peaks were called using macs2 version 2.1.2 (Zhang et al., 2008). Only peaks that were identified in all replicates for a given time point were used in downstream analyses. Peaks mapping to blacklist regions (Amemiya et al., 2019) and LINE/LTR repeat regions (Karolchik et al., 2004) were excluded from analyses. Peaks were assigned to genomic features and nearest genes using R package ChIPpeakAnno (Zhu et al., 2010). Bigwig tracks and line plots were generated using DeepTools utilities (Ramirez et al., 2014). Motif enrichment analysis was performed using homer version 4.11.1 (Heinz et al., 2010). Peak comparisons such as intersection with STARR-Seq enhancers were performed using bedtools utilities (Quinlan and Hall, 2010).

### RNA-Seq Data Analysis

Low-quality bases were trimmed using cutadapt 1.18 (Martin, 2011). Reads were aligned to DM6 or MM10 using STAR (Dobin et al., 2013). BAM files were generated using samtools 1.5 (Li et al., 2009). Read coverage per gene was calculated using featureCounts from subread version 1.6.0 (Liao et al., 2014). Normalized LogCPM values were generated in edgeR (Robinson et al., 2010). Bigwig tracks were generated using DeepTools utilities (Ramirez et al., 2014). Gene expression heatmaps were generated using R package pheatmap version 1.0.12 ._pheatmap: Pretty Heatmaps_. R package version 1.0.12, <https://CRAN.R-project.org/package=pheatmap>).

### ChIP-Seq Data Analysis

Reads were aligned to DM6 or MM10 using Bowtie2.4.1 (Langmead and Salzberg, 2012) using--local --very-sensitive parameters and max fragment size set to 1000bp. BAM files were generated using samtools 1.5 (Li et al., 2009). Bigwig tracks and line plots were generated using DeepTools utilities (Ramirez et al., 2014).

### Enhancer Reporter Assays

Gal4 driver lines were generated as part of the Janelia Flylight Gal4 or Vienna Tile collections (Kvon et al., 2014; Pfeiffer et al., 2008). These lines were crossed to G-TRACE reporter lines (Evans et al., 2009) and were reared and staged as described above. Samples were dissected and fixed as above, DAPI-stained, mounted in VectaShield, and imaged on a Leica DMI6000 or Leica SP5.

### HCR-FISH Assays

Detection of Stg and E2f1 transcripts *in situ* was performed using hybridization chain reaction (HCR)-fluorescent in situ hybridization (FISH) based on the protocol developed by Choi et al (Choi et al., 2018) and with adaptation to insects by Bruce and Patel (Bruce and Patel, 2020). Tissues were dissected in cold 1X PBS and fixed with 4% paraformaldehyde in 1X PBS for 30 minutes at room temperature (RT). Larval tissues were included in every dissection as a positive control for transcript expression. Fixed samples were washed 2 times in PBST (1X PBS with 0.1% Triton-X) for 5 minutes each. In the case of pupal wings, tissues were moved to a dissection dish for removal of the pupal cuticle surrounding the wings. Tissues were washed 2 times in 1X PBS for 5 minutes each, then moved to ice and washed in a series of increasing methanol solutions containing cold 25%, 50% and 75% methanol in 1X PBS for 5 minutes each. Tissues were washed with cold 100% methanol 2 times for 5 minutes each before storing in 100% methanol at −20C. To begin the hybridization protocol, tissues were rehydrated step-wise by moving to 75%, 50% and 25% methanol in PTw (1X PBS with 0.1% Tween-20) for 5 minutes each, then washed 2 times in PTw for 5 minutes each. Tissues were washed in Detergent Solution (1% SDS, 0.5% Tween-20, 50mM Tris-HCl, 1mM EDTA, 150mM NaCl) for 30 minutes at RT, then incubated in Probe Hybridization Buffer (Molecular Instruments, CA) at 37C for 30 minutes shaking at 600 rpm. Stg or E2f1 probe sets (Molecular Instruments) were used overnight at 4nM in Probe Hybridization Buffer at 37C and shaking at 600 rpm. Samples were washed with Probe Wash Buffer (Molecular Instruments) at 37C, shaking at 600 rpm, 4 times for 15 minutes each. Tissues were then washed with 5X SSCT (5X SSC buffer and 0.1% Tween-20) 2 times at RT for 5 minutes each. Samples were incubated in Amplification Buffer (Molecular Instruments) at RT for 30 minutes. Fluorescent hairpins (Molecular Instruments) were prepared by heating at 95C for 90 seconds, then cooling at RT for 30 minutes in the dark. Hairpins were applied to samples at 60nM in Amplification Buffer and kept overnight at 23C with shaking at 600 rpm, and protected from light from this point forward. Samples were washed with 5X SSCT at RT 2 times for 5 minutes each, then 2 times for 30 minutes each, then 1 time for 5 minutes. Tissues were stained with 1ng/mL DAPI in PBST for 10 minutes prior to mounting in Vectashield and imaging on a Leica SP8 confocal microscope.

### Data Access

Data generated in this study can be accessed from GEO GSE263160. Previously published datasets: Larval and pupal wing ATAC-Seq data can be accessed from GEO GSE211152. Larval and pupal wing RNA-Seq data can be accessed from GEO GSE131981. ENCODE-generated RNA-Seq data from larval and 2 day pupa brain can be accessed from NCBI BioProject PRJNA75285, samples SRX029398, SRX042030, and SRX029404. STARR-Seq data can be accessed from GEO GSE57876. Trl ChIP-Seq from wing disc can be accessed from GEO GSE38594. DREF ChIP-Seq from Kc cells can be accessed from GEO GSE39664. M1BP ChIP-Seq from Kc cells can be accessed from GEO GSE142531. Myc ChIP-Seq from Kc cells can be accessed from GEO GSE39521. RNA-Seq, ATAC-Seq, and ChIP-Seq datasets from developing mouse retina can be accessed from GEO GSE87064.

## Acknowledgements

We thank Dr. Erick Bayala Rodriguez for assistance with HCR-FISH. This work was supported by NIH/NIGMS (R01) GM127367 and NIH/NIGMS (R35) GM149273 to LB. EMB was supported by the U. Michigan Genetics Training Grant (T32GM007544). We thank members of the Buttitta Lab for advice and feedback on this work.

