## Supplement for "Transcriptional repression and enhancer decommissioning silence cell cycle genes in postmitotic tissues"

**Contains Supplemental Figures 1-13 with legends**

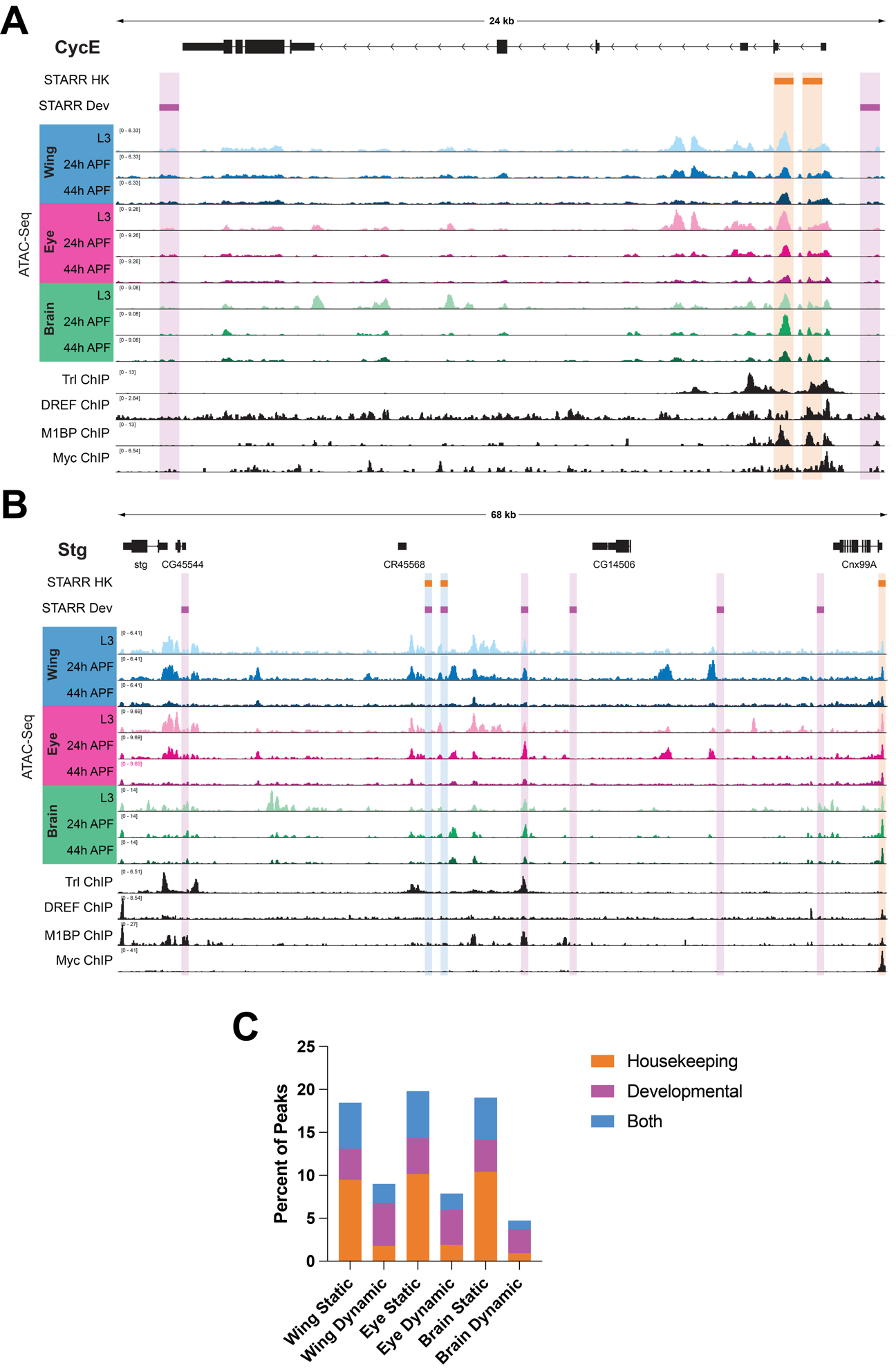

**Supplemental Fig. 1**

**STARR-Seq fails to identify many developmentally dynamic enhancer regions *in vivo.*** (**A-B**) STARR-Seq, ATAC-Seq, and ChIP-Seq data at the *CycE* (**A**) and *stg* (**B**) loci. Orange boxes indicate STARR-Seq housekeeping enhancers and magenta boxes indicate STARR-Seq developmental enhancers. ATAC-Seq accessibility data from wing, eye, and brain at L3, 24h APF, and 44h APF. ChIP-Seq data for Trl, DREF, M1BP, and Myc. Y-axes indicate the normalized read counts per million. HK, housekeeping. Dev, developmental. (**C**) Bar plots indicating the percent of ATAC-Seq peaks that overlapped housekeeping, developmental, or both types of STARR-Seq defined enhancers, grouped by those that showed statistically significant changes in ATAC-Seq intensity across the time course (Dynamic) versus those that did not (Static).

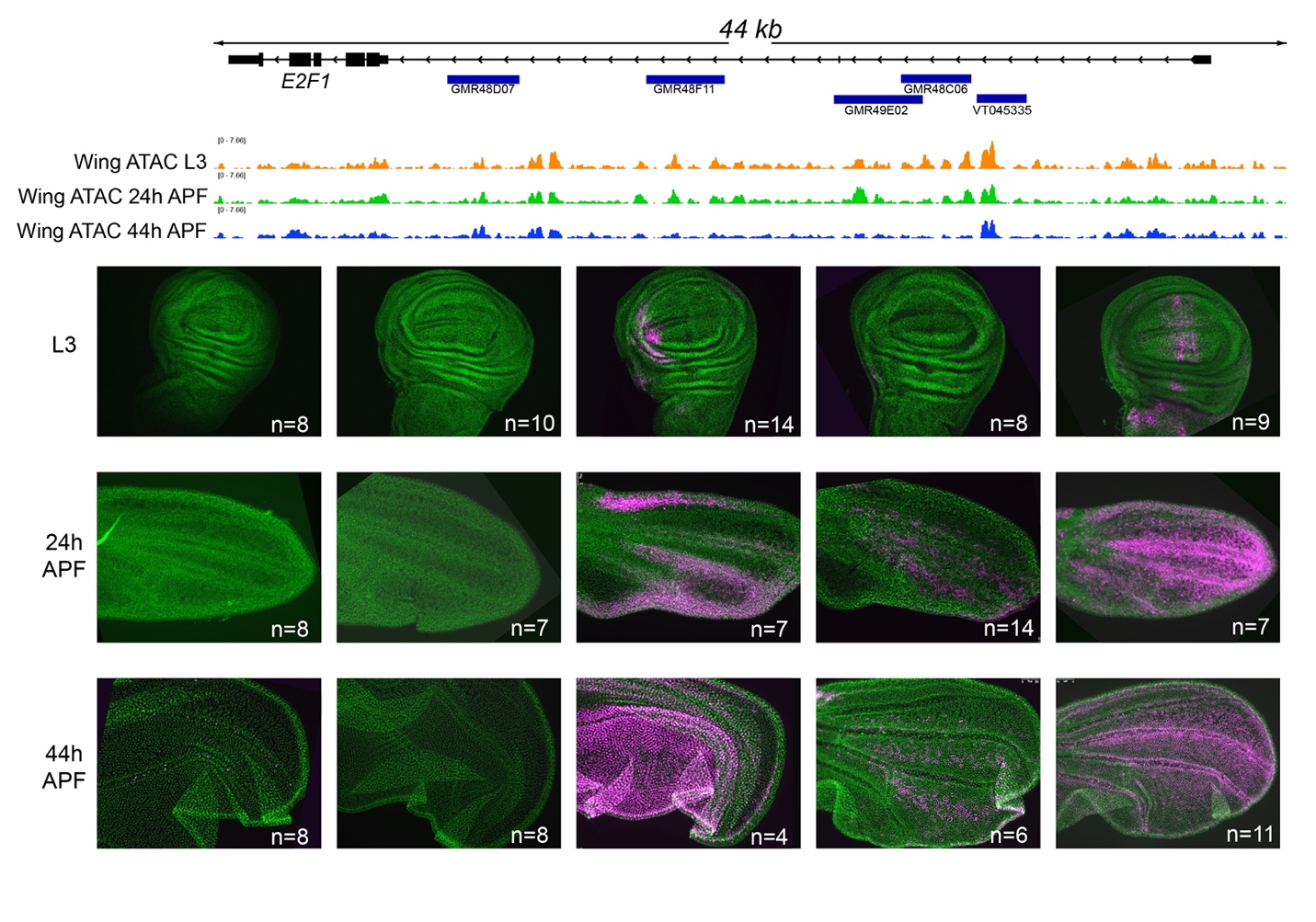

**Supplemental Fig. 2**

**Dynamic chromatin regions within *e2f1* function together to drive expression of *e2f1* transcript throughout the wing.** ATAC-seq shows chromatin accessibility in the wing during development. We observe that a few of these regions show enhancer activity in the wing. These enhancers may function in a modular fashion, as the sum of their expression patterns drive expression of *e2f1* throughout the pupal wing. The indicated Gal4 lines were crossed to G-TRACE with UAS-RFP (current expression) in magenta. DNA was stained with DAPI and shown in green. Note that intensity has been adjusted across each image to emphasize current expression. For intensity quantifications in Supp. Figs. 8 and 9 images were acquired under identical settings and intensity adjustments were not performed.

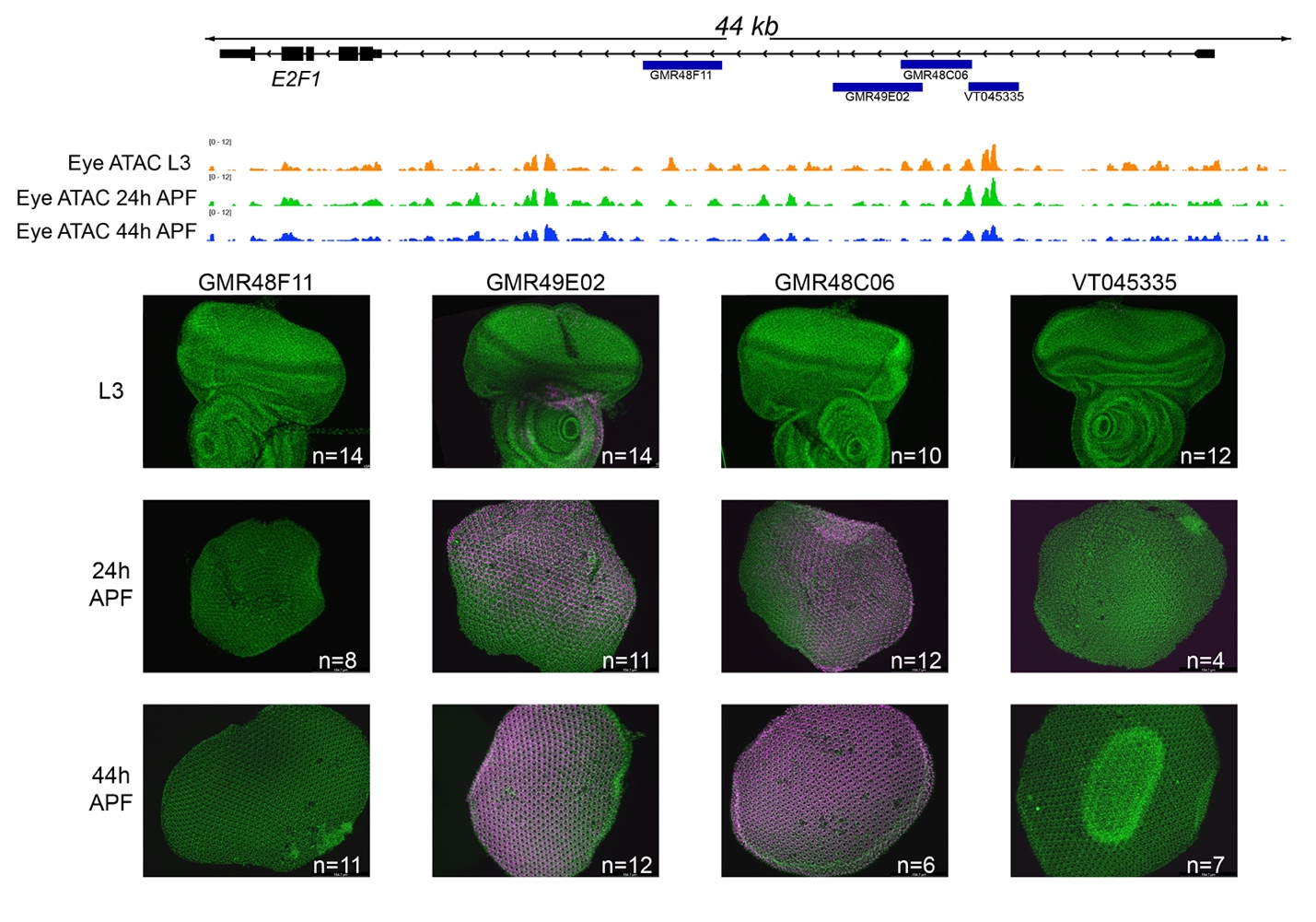

**Supplemental Fig. 3 Dynamic chromatin regions within *e2f1* show enhancer activity in the pupal eye.**

ATAC-seq data shows chromatin accessibility changes throughout development. The indicated Gal4 lines were crossed to G-TRACE with UAS-RFP (current expression) in magenta. DNA was stained with DAPI and shown in green. Note that intensity has been adjusted equally across entire images to emphasize current expression. For intensity quantifications in Supp. Figs. 8 and 9 images were acquired under identical settings and intensity adjustments were not performed.

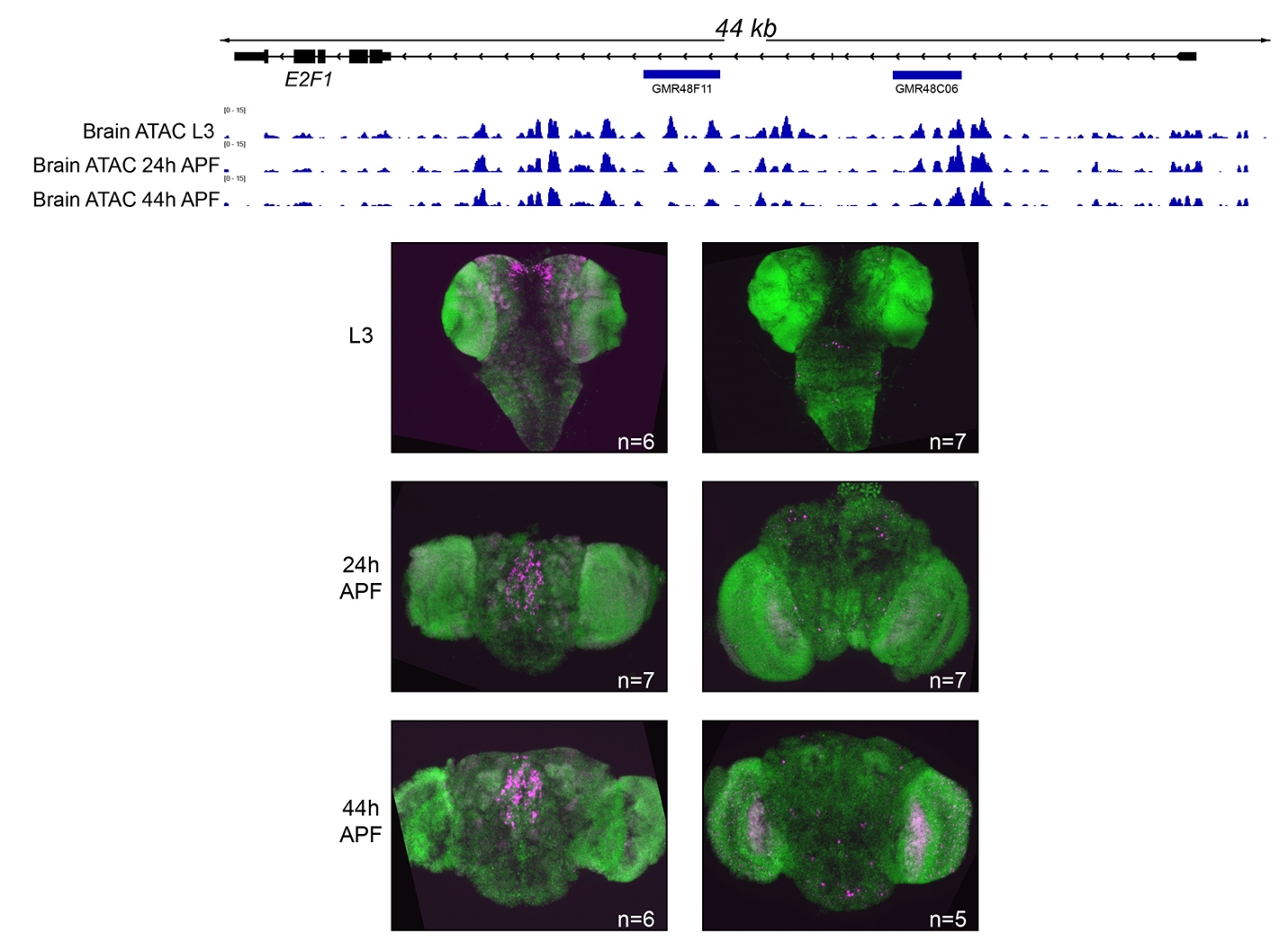

**Supplemental Fig. 4**

**Dynamic chromatin regions within *e2f1* show enhancer activity in the brain.**

ATAC-seq data shows chromatin accessibility changes throughout development. The indicated Gal4 lines were crossed to G-TRACE with UAS-RFP (current expression) in magenta. DNA was stained with DAPI and shown in green. Note that intensity has been adjusted equally across entire images to emphasize current expression. For intensity quantifications in Supp. Figs. 8 and 9 images were acquired under identical settings and intensity adjustments were not performed.

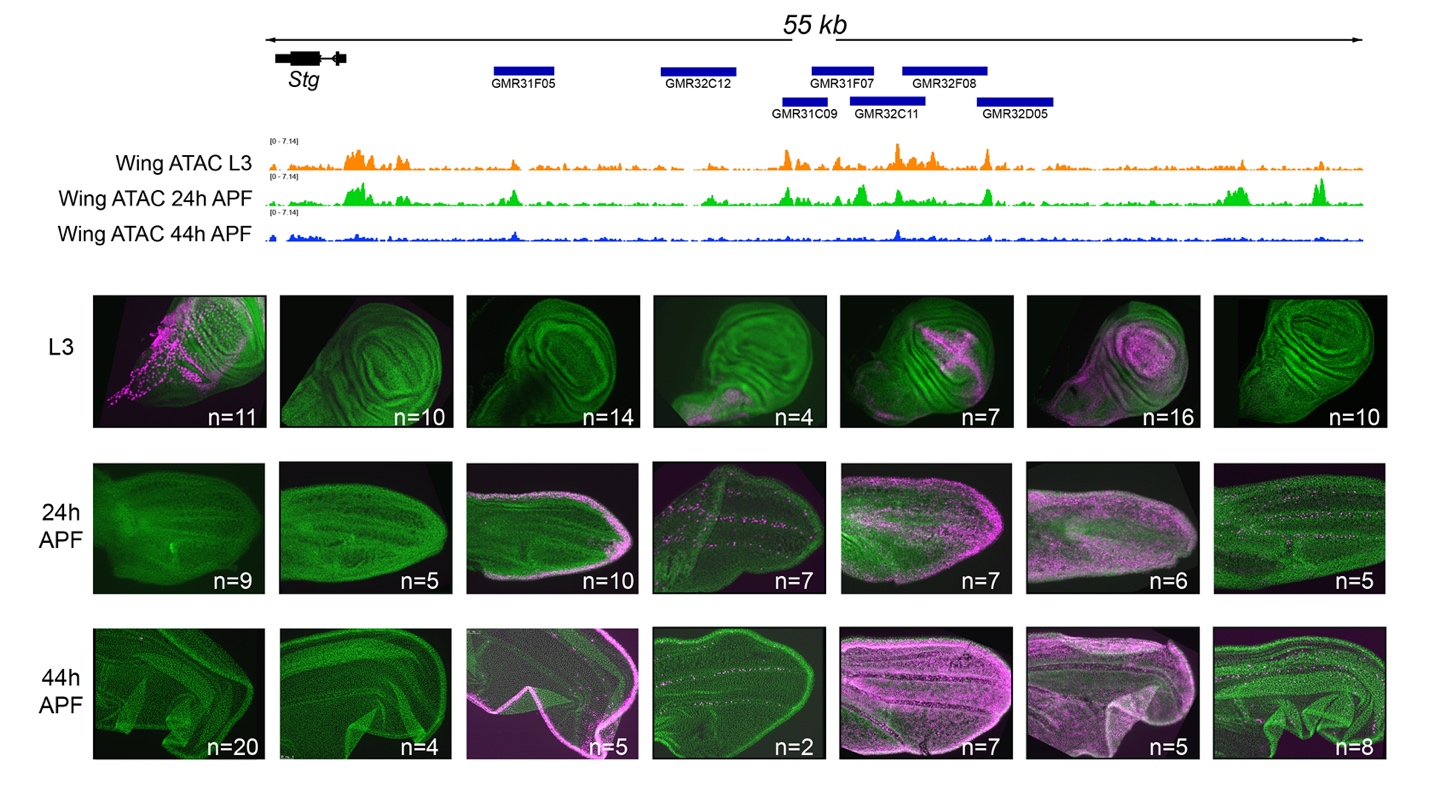

**Supplemental Fig. 5 Modular enhancers function together to drive *stg* expression**

**throughout the larval and pupal wing.** ATAC-seq data shows dynamic chromatin accessibility changes throughout metamorphosis. The indicated Gal4 lines were crossed to G-TRACE with UAS-RFP (current expression) in magenta. DNA was stained with DAPI and shown in green. Note that intensity has been adjusted equally across entire images to emphasize current expression. For intensity quantifications in Supp. Figs. 8 and 9 images were acquired under identical settings and intensity adjustments were not performed.

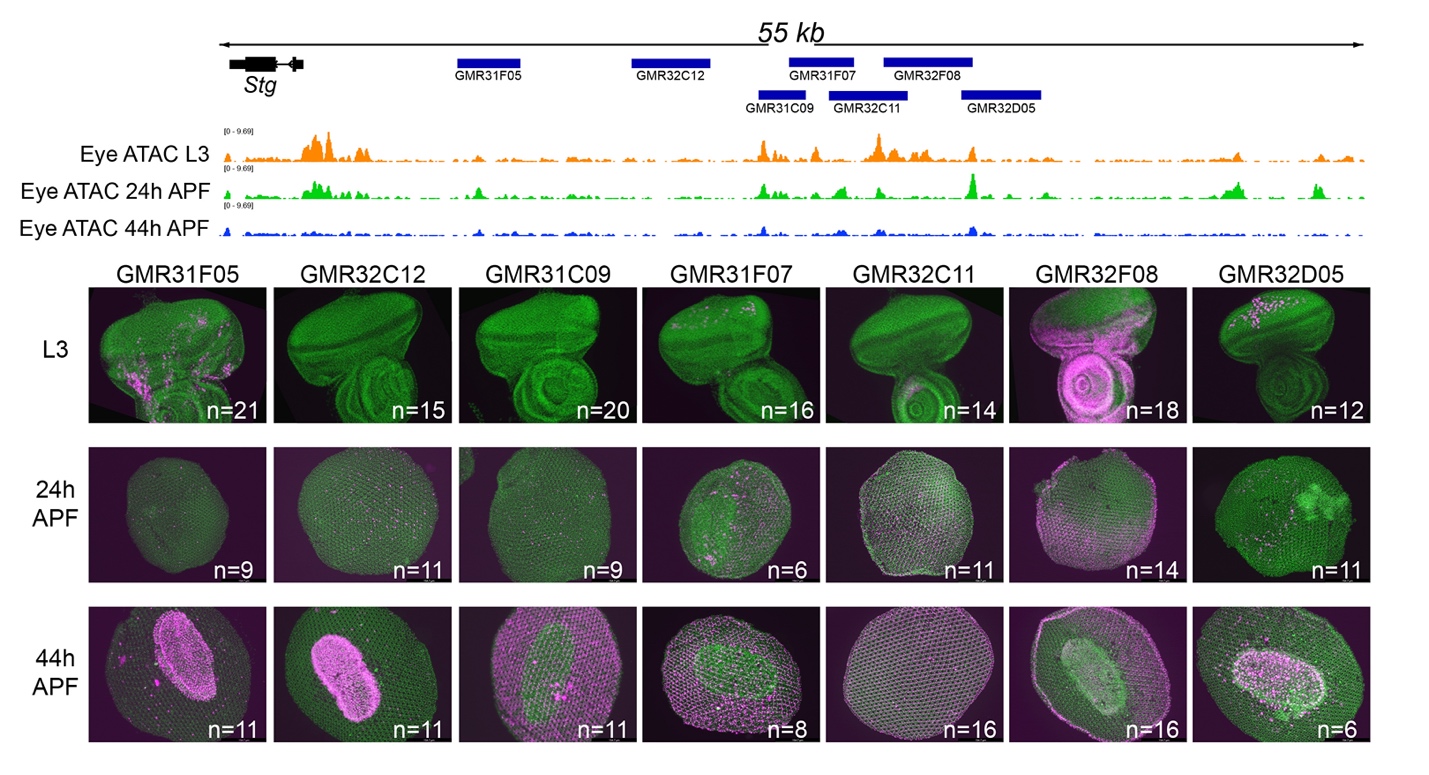

**Supplemental Fig. 6 Dynamic chromatin regions in the *stg* cis-regulatory region have**

**enhancer activity in the larval and pupal eye**.

Eye ATAC-seq data shows accessible chromatin throughout metamorphosis. The indicated Gal4 lines were crossed to G-TRACE with UAS-RFP (current expression) in magenta. DNA was stained with DAPI and shown in green. Note that intensity has been adjusted equally across entire images to emphasize current expression. For intensity quantifications in Supp. Figs. 8 and 9 images were acquired under identical settings and intensity adjustments were not performed.

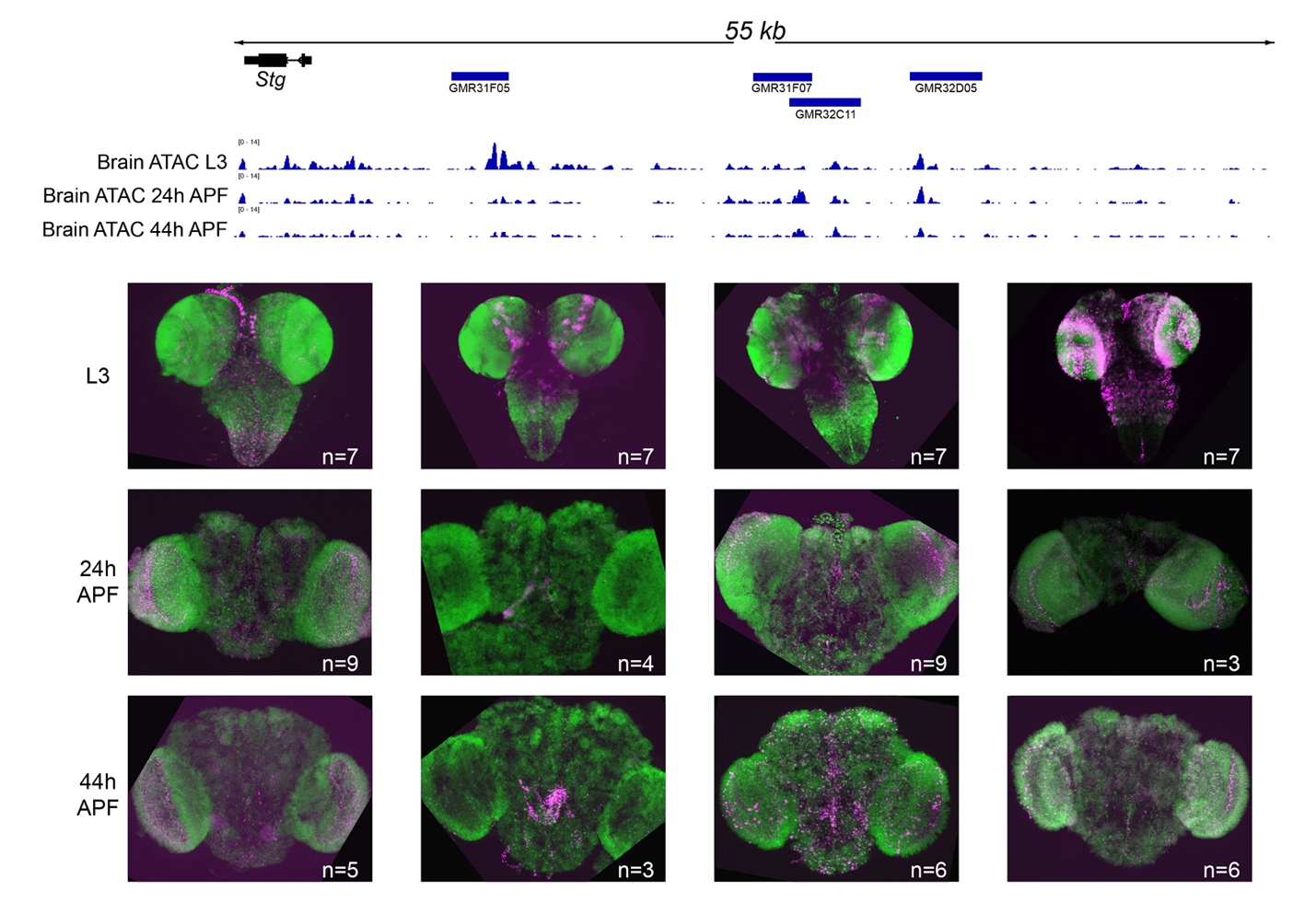

**Supplemental Fig. 7 Dynamic chromatin regions in the *stg* cis-regulatory region have**

**enhancer activity in the larval and pupal brain.**

The indicated Gal4 lines were crossed to G-TRACE with UAS-RFP (current expression) in magenta. DNA was stained with DAPI and shown in green. Note that intensity has been adjusted equally across entire images to emphasize current expression. For intensity quantifications in Supp. Figs. 8 and 9 images were acquired under identical settings and intensity adjustments were not performed.

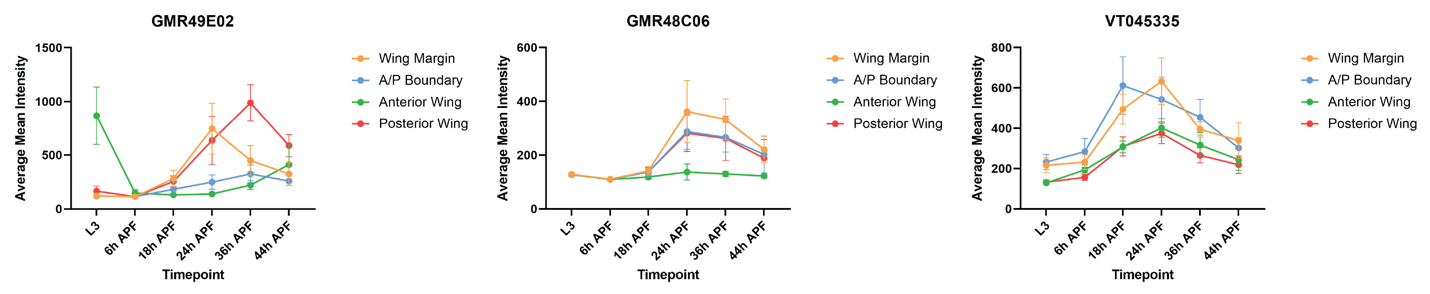

**Supplemental Fig. 8**

Quantifications of enhancer activity for the indicated *e2f1* Gal4 lines crossed to G-Trace in the across wings. The indicated Gal4 lines were crossed to G-TRACE with UAS-RFP (current expression) quantified. Images were acquired under identical settings without intensity adjustments. Quantifications were performed using Integrated intensity normalized to background in Image J.

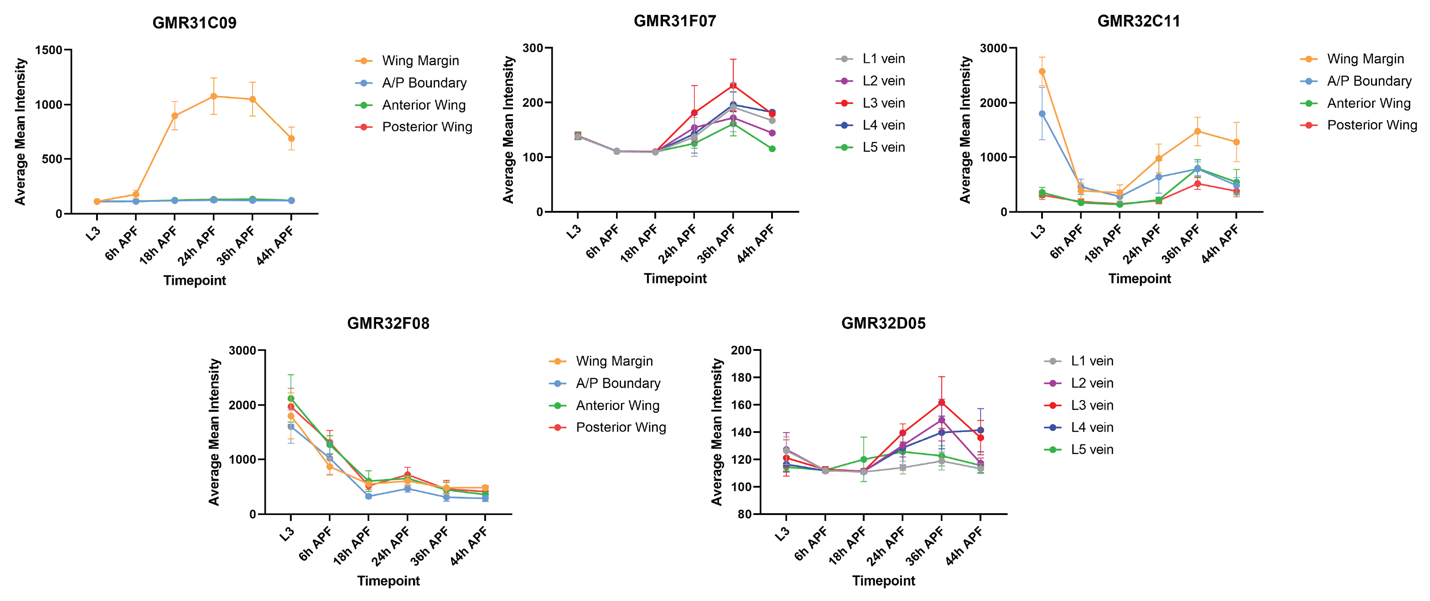

**Supplemental Fig. 9**

Quantifications of enhancer activity for the indicated *stg* Gal4 lines crossed to G-Trace in the across wings. The indicated Gal4 lines were crossed to G-TRACE with UAS-RFP (current expression) quantified. Images were acquired under identical settings without intensity adjustments. Quantifications were performed using Integrated intensity normalized to background in Image J.

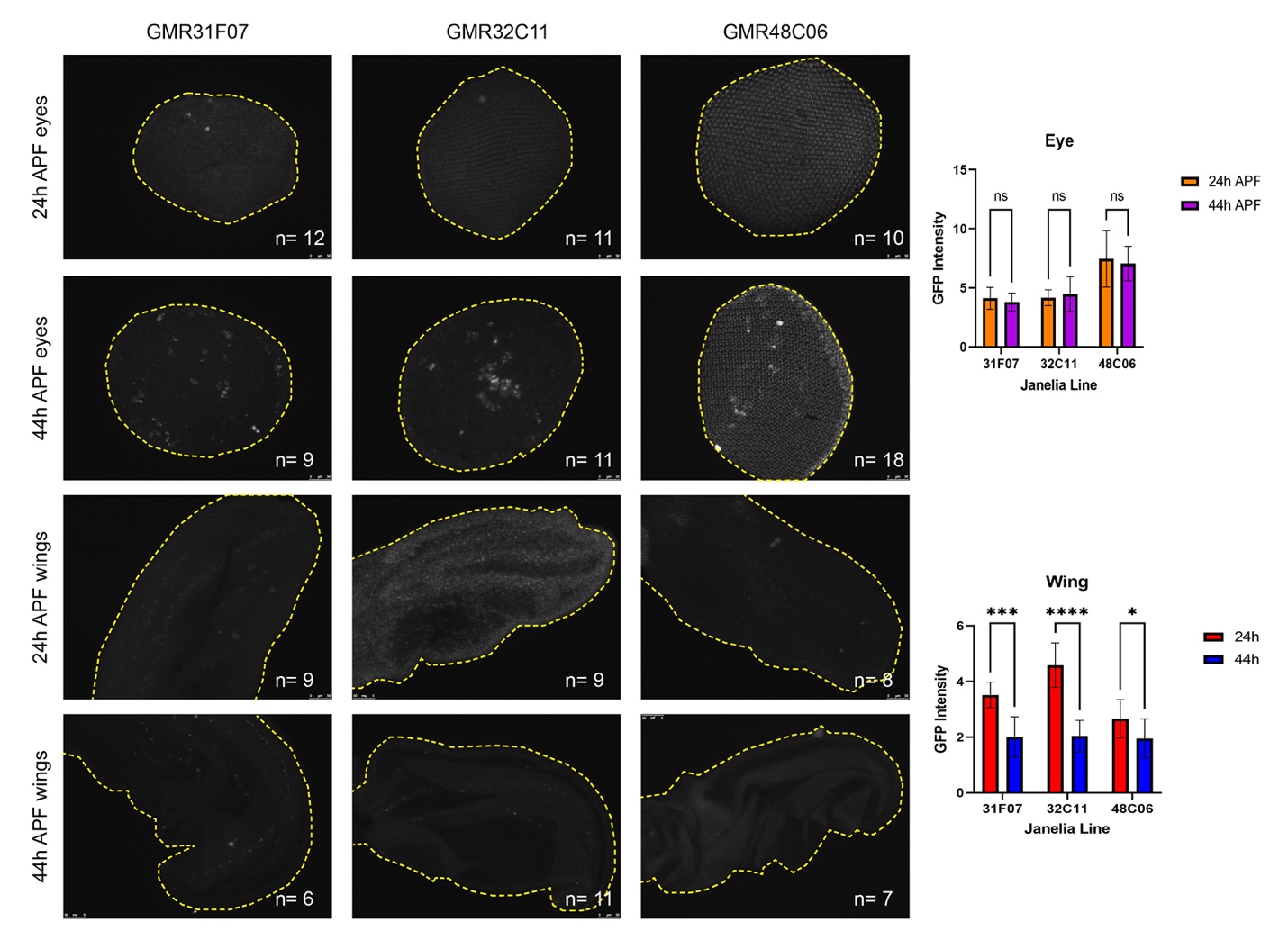

**Supplemental Fig. 10**

The indicated Gal4 lines were crossed to UAS-de-stabilized GFP (ds-GFP) in grey. Tissue outlines are shown with yellow dotted lines. Quantifications of enhancer activity using dsGFP expression are shown. Images were acquired under identical settings without intensity adjustments. Quantifications were performed using Integrated intensity normalized to background in Image J.

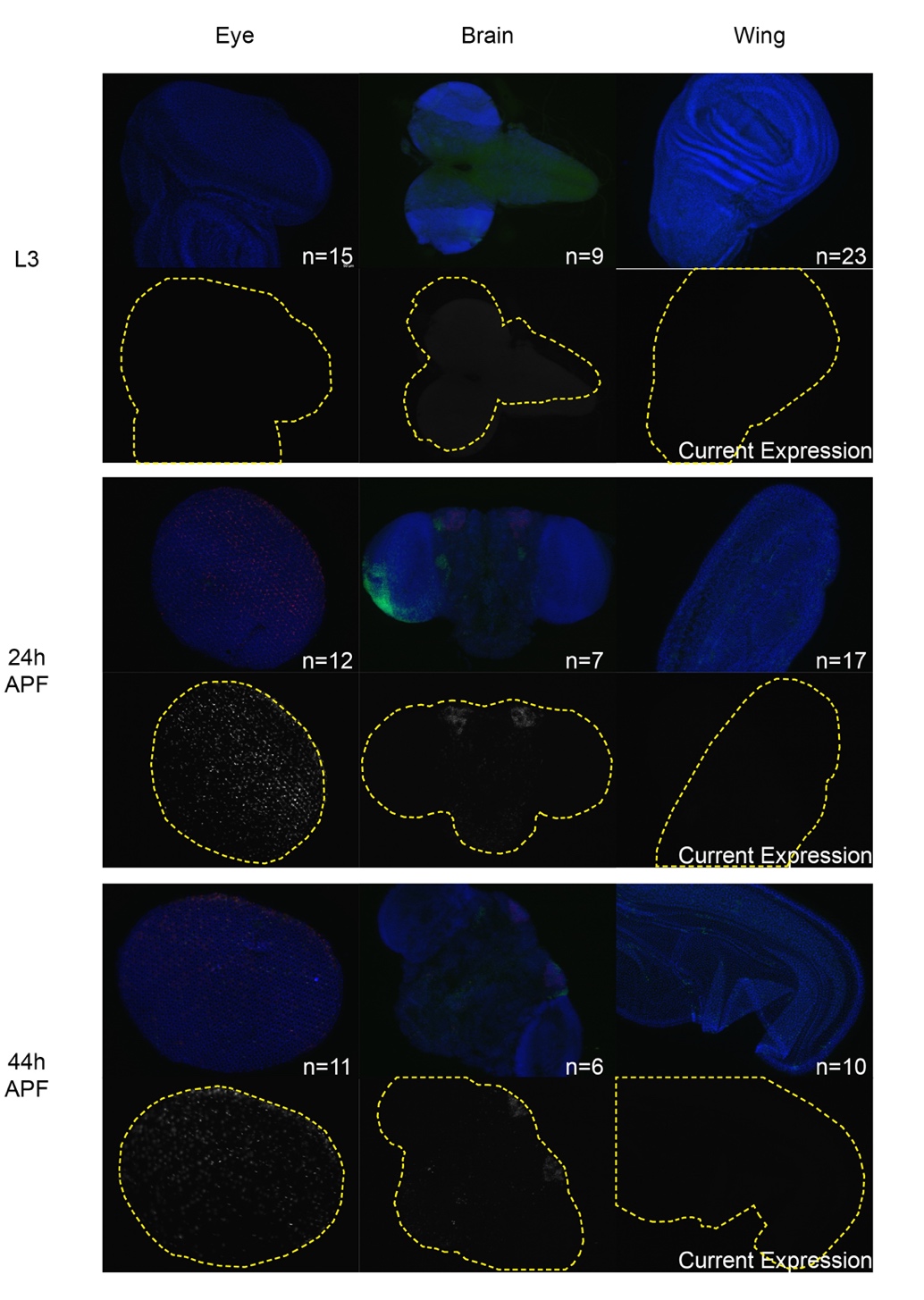

**Supplemental Fig. 11 ‘Empty’ Janelia-Gal4 line shows some current expression in pupal eyes and brains.**

The control Janelia Gal4 line without an inserted genomic fragment, containing the Drosophila synthetic Core promoter (DSCP) was crossed to G-TRACE with UAS-RFP (current expression) in grey. Past lineage-tracing expression is in green and DNA was stained with DAPI and shown in blue. Note that intensity has been adjusted equally across entire images to emphasize current expression. We note significant expression from the DSCP in pupal eyes and brains, which may confound enhancer quantifications at 24 and 44h APF.

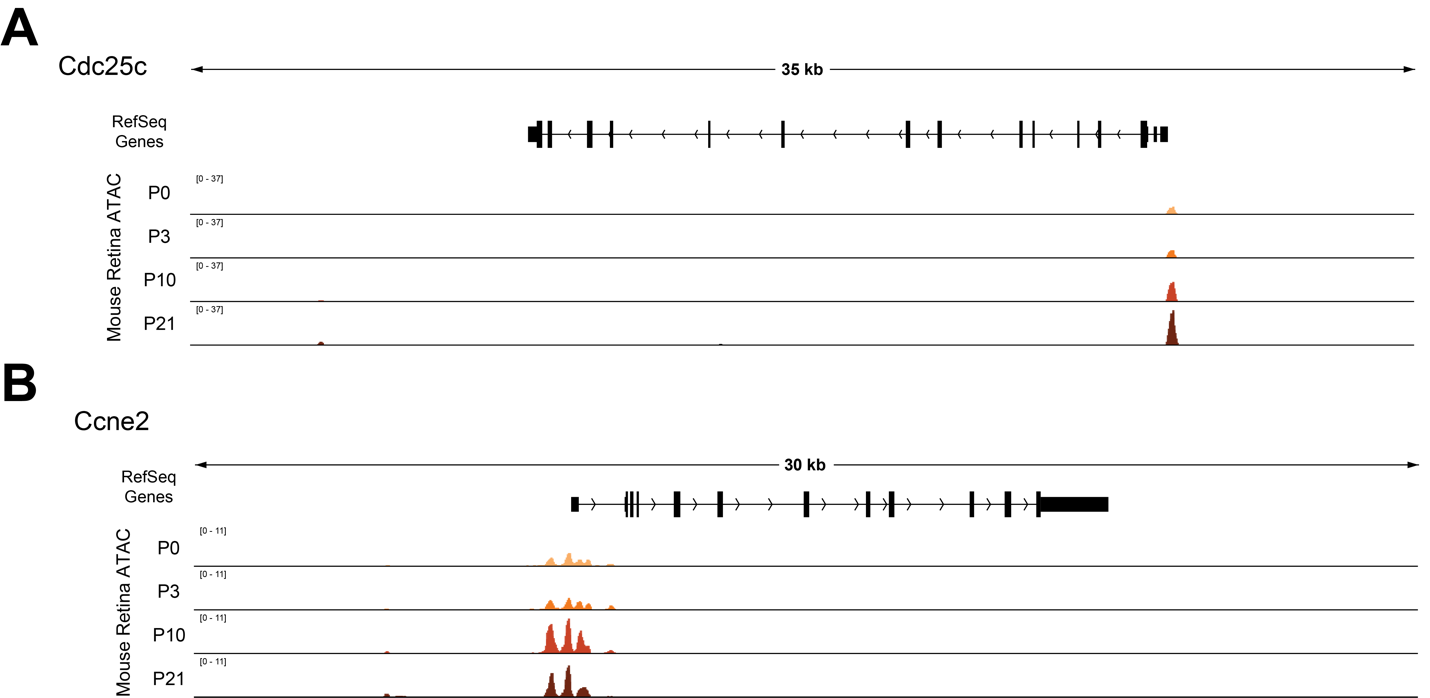

**Supplemental Fig. 12**

**Loci encoding the orthologs of Stg and CycE provide no obvious indication of postmitotic enhancer decommissioning in the developing mouse retina.** ATAC-Seq data tracks from the mouse retina at the *Cdc25c* locus (**A**, ortholog of Stg in flies) and the Ccne2 locus (**B**, ortholog of CycE in flies).

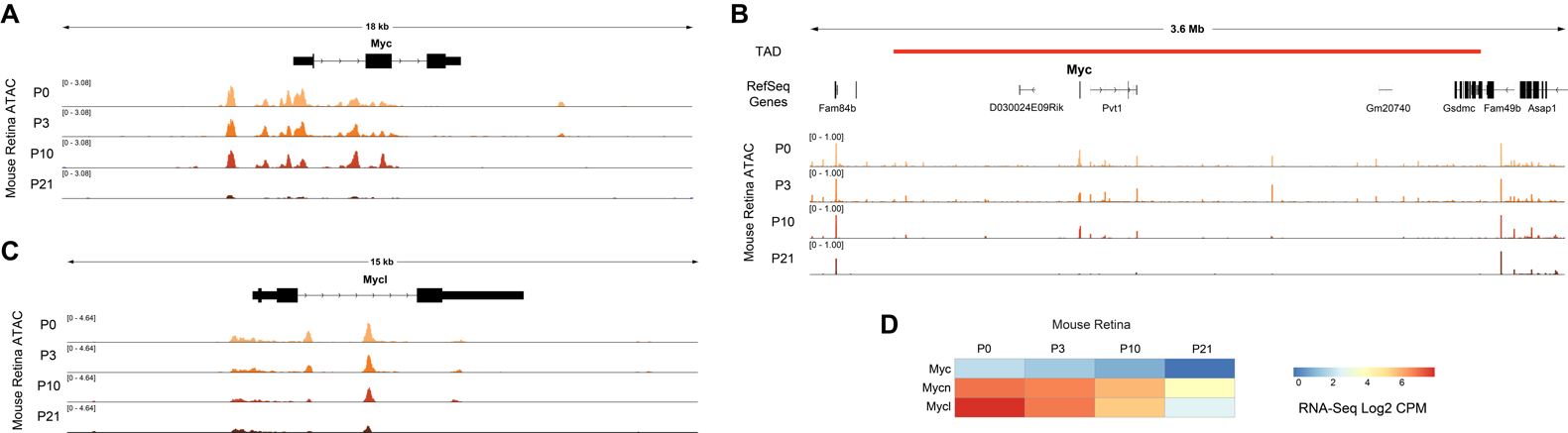

**Supplemental Fig. 13**

**Loci encoding the Myc family members provide evidence of postmitotic enhancer decommissioning in the developing mouse retina.** (**A-C**) ATAC-Seq data tracks from the mouse retina at the *Myc* gene body (**A**), at the topologically associating domain (TAD) containing *Myc* (**B**), and the *Mycl* locus (**C**). (**D**) Heatmap depicting the average transcript expression values for Myc family genes. Data are presented as normalized Log2 Count Per Million (CPM) values.
